# A role for the Gram-negative outer membrane in bacterial shape determination

**DOI:** 10.1101/2023.02.03.527047

**Authors:** Elayne M. Fivenson, Patricia D.A. Rohs, Andrea Vettiger, Marios F. Sardis, Grasiela Torres, Alison Forchoh, Thomas G. Bernhardt

**Affiliations:** Department of Microbiology, Blavatnik Institute, Harvard Medical School, Boston, MA 02115; Howard Hughes Medical Institute, Boston, United States

**Keywords:** peptidoglycan, lipopolysaccharide, morphogenesis, cell envelope

## Abstract

The cell envelope of Gram-negative bacteria consists of three distinct layers: the cytoplasmic membrane, a cell wall made of peptidoglycan (PG), and an asymmetric outer membrane (OM) composed of phospholipid in the inner leaflet and lipopolysaccharide (LPS) glycolipid in the outer leaflet. The PG layer has long been thought to be the major structural component of the envelope protecting cells from osmotic lysis and providing them with their characteristic shape. In recent years, the OM has also been shown to be a load-bearing layer of the cell surface that fortifies cells against internal turgor pressure. However, whether the OM also plays a role in morphogenesis has remained unclear. Here, we report that changes in LPS synthesis or modification predicted to strengthen the OM can suppress the growth and shape defects of *Escherichia coli* mutants with reduced activity in a conserved PG synthesis machine called the Rod system (elongasome) that is responsible for cell elongation and shape determination. Evidence is presented that OM fortification in the shape mutants restores the ability of MreB cytoskeletal filaments to properly orient the synthesis of new cell wall material by the Rod system. Our results are therefore consistent with a role for the OM in the propagation of rod shape during growth in addition to its well-known function as a diffusion barrier promoting the intrinsic antibiotic resistance of Gram-negative bacteria.

**SIGNIFICANCE:** The cell wall has traditionally been thought to be the main structural determinant of the bacterial cell envelope that resists internal turgor and determines cell shape. However, the outer membrane (OM) has recently been shown to contribute to the mechanical strength of Gram-negative bacterial envelopes. Here, we demonstrate that changes to OM composition predicted to increase its load bearing capacity rescue the growth and shape defects of *Escherichia coli* mutants defective in the major cell wall synthesis machinery that determines rod shape. Our results therefore reveal a previously unappreciated role for the OM in bacterial shape determination in addition to its well-known function as a diffusion barrier that protects Gram-negative bacteria from external insults like antibiotics.

## INTRODUCTION

Gram-negative bacteria have a characteristic three-layered cell envelope comprised of an inner (cytoplasmic) membrane (IM), a relatively thin cell wall made of peptidoglycan (PG), and an outer membrane (OM). The OM bilayer is asymmetric with phospholipids in the inner leaflet and the lipopolysaccharide (LPS) glycolipid in the outer leaflet. For many years, the PG layer was thought to be the sole load-bearing component of the envelope with the OM primarily serving to protect Gram-negative cells from external insults like antibiotics (1,2). However, it has recently become clear that in addition to providing a barrier function, the OM can also help cells resist internal turgor pressure (3). What has remained unknown is whether the OM also partners with the PG layer to define cell shape. Here, we report a genetic analysis of PG synthesis and cell shape determination that supports such a role for the OM.

The PG heteropolymer is composed of glycan chains with alternating units of N-acetylglucosamine (GlcNAc) and N-acetylmuramic acid (MurNAc) (4). A short peptide is attached to the MurNAc sugar and is used to crosslink adjacent glycans to form the cell wall matrix. Glycosyltransferases (GTases) catalyze the polymerization of glycan polymers whereas transpeptidases (TPases) perform the crosslinking reaction. There are two major classes of PG synthases: class A Penicillin Binding Proteins (aPBPs) and complexes formed between SEDS (Shape, Elongation, Division, Sporulation) proteins and class B PBPs (bPBPs) (1,2, 5). The aPBPs have both enzymatic functions in a single polypeptide whereas in the SEDS-bPBP complexes, the SEDS protein promotes glycan polymerization and the bPBP provides the crosslinking activity (6–9).

The SEDS-bPBP complexes RodA-PBP2 (6–8, 10) and FtsW-FtsI (9) play essential roles in rod shape determination and cell division, respectively. In both cases, these synthases are part of larger multiprotein assemblies involving cytoskeletal filaments. The rod shape determining system is called the Rod system (a.k.a. the elongasome). It promotes the elongation of bacilli and maintains their characteristic rod shape. In addition to RodA-PBP2, the system includes filaments of the actin-like MreB protein along with three membrane proteins of poorly understood function: MreC, MreD, and RodZ (11–18). The Rod system has been observed to dynamically rotate around the long axis of the cell as it deposits new PG material to promote cell elongation. PG synthesis is required for the motion and MreB filaments are thought to orient it orthogonally to the long cell axis via a rudder-like mechanism (1, 7, 19–22).

To better understand Rod system function, we previously identified non-functional variants of MreC in *Escherichia coli* and selected for suppressor mutations that overcame their shape and viability defects (10, 23). One major class of suppressors encoded hypermorphic variants of PBP2 and RodA that provided important insight into the Rod system activation mechanism and the regulation of SEDS proteins (10). Genetic, structural, and cytological evidence suggests that MreC activates the system by inducing a conformational change in PBP2, which in turn activates RodA, shifting the complex from an inactive to an active state (10). The role of MreD in the complex is not clear (23, 24). The signals that promote Rod system activation also remain unknown, but the mechanism may involve the recognition of landmarks in the PG matrix by PBP2 (25).

In this report, we study a new class of suppressors that restore the growth and shape of *mreC* hypomorphs. Instead of activating the Rod system directly, these suppressors function by increasing the production of LPS. Further analysis of the suppression mechanism revealed that Rod system mutants are impaired for LPS production. Additionally, we found that modifications to LPS predicted to stiffen the OM restore rod shape in cells defective for MreC by promoting the feedback mechanism via which MreB orients PG synthesis. Thus, our results suggest a potential connection between Rod system activity and LPS synthesis and argue for a morphogenic role for the OM.

## RESULTS

### Increased LPS synthesis suppresses a Rod system defect

Cells with *mreC(R292H*) or *mreC(G156D*) mutations produce stable MreC protein capable of inducing a dominant-negative growth and shape phenotype (10, 23). Therefore, the altered proteins are likely capable of joining the Rod complex but are defective in stimulating its activity. Mutants with these alleles at the native locus can be maintained as spheres on minimal medium (M9), but they fail to grow on rich medium (LB). We selected for spontaneous suppressors that restored the growth of these mutants on LB along with their rod shape. In addition to mutants encoding altered PBP2 and RodA described previously (10), the selection also identified suppressors in the *ftsH* and *lapB(yciM*) genes encoding regulators of LPS synthesis (**Fig. 1, SI Table 1**). FtsH is an IM metalloprotease that along with its adapter protein LapB (26, 27) degrades LpxC (UDP-3-O-acyl-N-acetylglucosamine deacetylase) (28–30), the enzyme that catalyzes the first committed step in LPS synthesis (31, 32). Proteolysis of LpxC is in turn regulated by the essential inner membrane protein YejM (PbgA, LapC), which functions to inhibit LapB activity in a manner that is sensitive to the concentration of LPS in the IM (30, 33–38). When the steady state concentration is low due to LPS synthesis being balanced with its transport to the OM, YejM blocks LpxC turnover (**Fig. 1A, top**). However, when LPS synthesis outpaces its transport, YejM is inhibited by the buildup of LPS in the inner membrane and LpxC turnover is increased to restore homeostasis (**Fig. 1A, bottom**).

**Figure 1:**
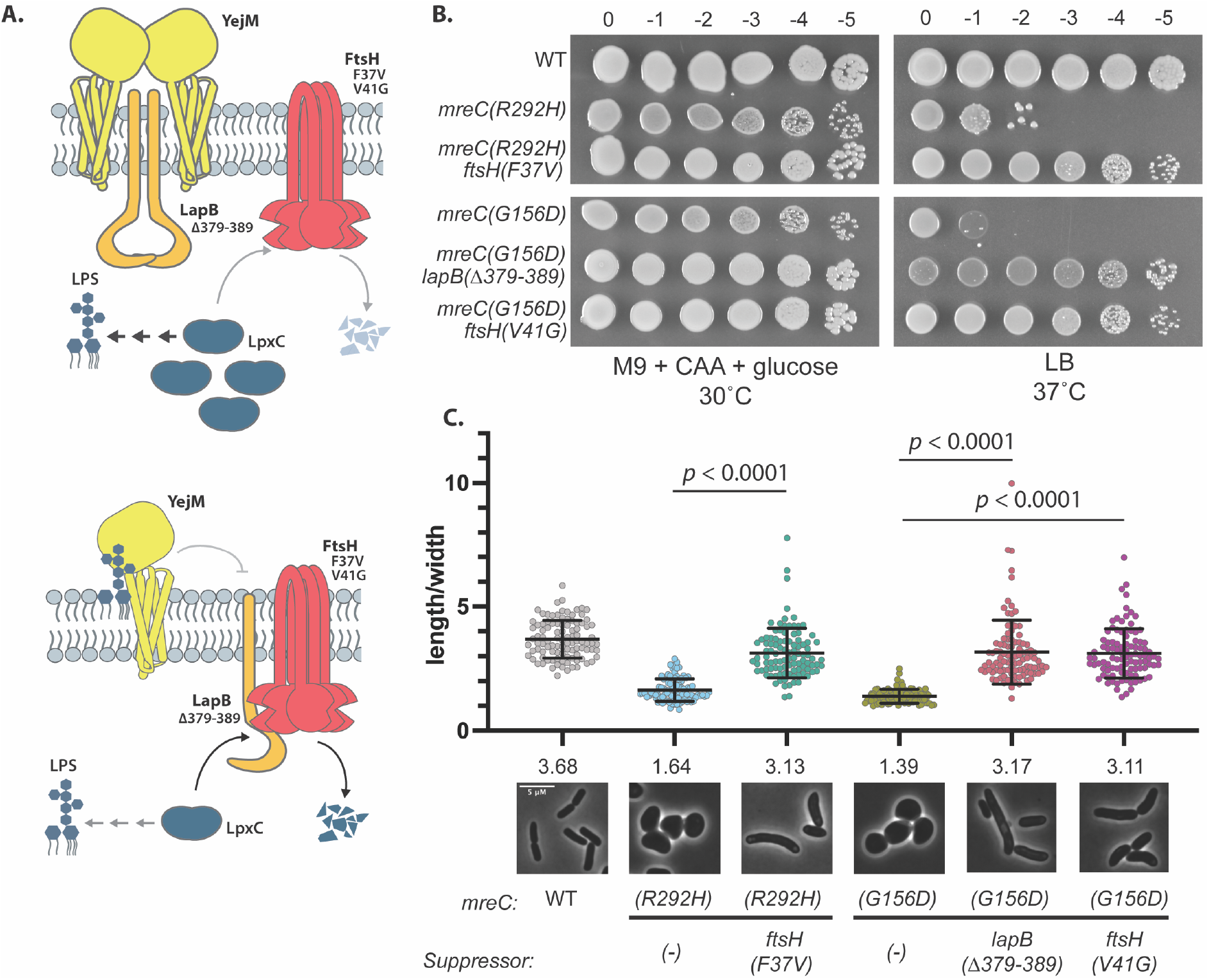
Mutations in factors involved in LpxC turnover rescue *mreC* hypomorphs. **A**. Schematic overview of LpxC regulation (suppressors mutations are noted below protein names). TOP: When LPS levels are low, YejM interacts with LapB, sequestering it from the FtsH protease, leading to the stabilization of LpxC and increased LPS synthesis. BOTTOM: When LPS levels are high, LPS accumulates in the outer leaflet of the inner membrane. YejM binds to LPS, allowing LapB to interact with FtsH and target LpxC for degradation, reducing LPS synthesis. **B**. WT (HC555), *mreC(R292H*) (PR5), *mreC(R292H) ftsH(V37G*) (PR82), *mreC(G156D*) (PR30), *mreC(G156D) lapB(Δ379-389*) (PR86), *mreC(G156D) ftsH(V41G*) (PR88) were cultured for 24 hours in minimal medium (M9 + CAA + glu) at 30°C. Cultures were then normalized to OD_600_= 1 and serially diluted and spotted onto LB and M9 + CAA + glu plates. LB plates were incubated for 16 hours at 37°C and M9 plates were incubated for 40 hours at 30°C. Dilution factors are indicated above the spot dilutions. **C**. Micrographs of WT (HC555), *mreC(R292H*) (PR5), *mreC(R292H) ftsH(V37G*) (PR82), *mreC(G156D*) (PR30), *mreC(G156D) lapB(Δ379-389*) (PR86), *mreC(G156D) ftsH(V41G*) (PR88). Strains were grown overnight in minimal medium (M9 + CAA + glu) at 30°C. Overnight cultures were then back diluted to OD_600_= 0.05 in minimal medium and incubated shaking at 30°C until OD_600_= 0.3-0.4. Cells were then spun down and resuspended in LB to an OD_600_ of 0.025 and incubated at 37°C until OD_600_=0.3-0.4. Cells were then fixed and imaged. Aspect ratios were analyzed using the FIJI plugin MicrobeJ (70). Scale bar = 5 μm. n= 100 cells per group. Statistical significance determined using an Unpaired t test with Welch’s correction (not assuming equal SDs).

Both *ftsH* suppressors encoded protease variants with substitutions in the periplasmic loop of the protein (**Fig. 1**). One was found as a suppressor of *mreC(R292H*) and the other as a suppressor of *mreC(G156D*) (**Fig. 1, SI Table 1**). A mutation in *lapB* encoding a protein with a deletion of the last eleven C-terminal amino acids was also isolated as a suppressor of *mreC(G156D*) (**Fig. 1, SI Table 1**). Although growth rate and morphology were not restored to completely match those of wild-type cells, the suppressors supported full plating efficiency of their respective *mreC* mutant on LB (**Fig. 1B**) and switched their morphology from sphere-like to elongated rods (**Fig. 1C**). Suppression was not allele specific as the *ftsH(V41G*) mutation originally isolated as a suppressor of *mreC(G156D*) (**SI Table 1**) also suppressed the growth and shape defects of *mreC(R292H*) (**Fig. 2A-B**).

**Figure 2:**
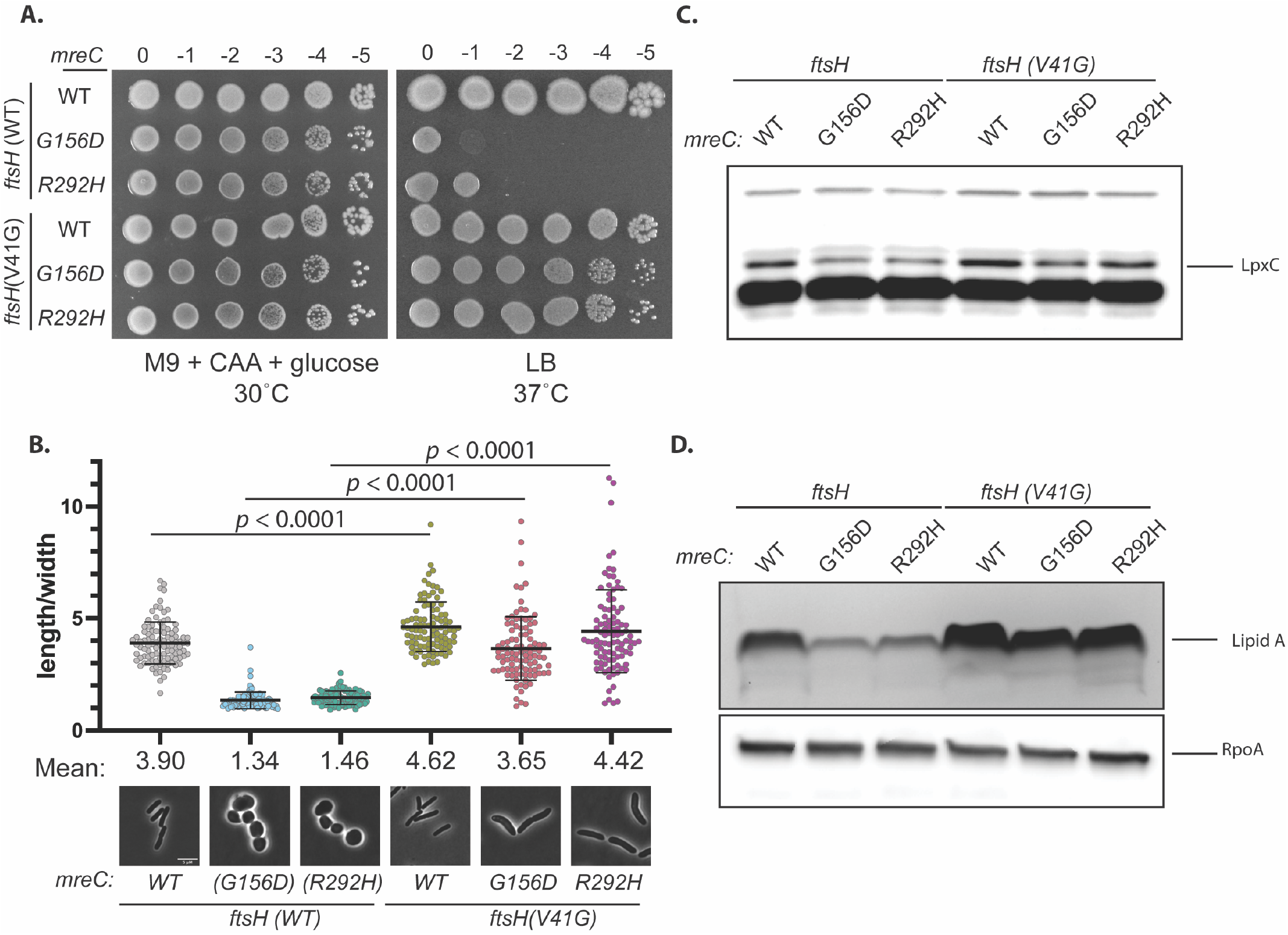
*FtsH(V41G*) increases LpxC and LPS levels in *mreC* hypomorphs. **A.**Cultures of WT(EMF196), *mreC(G156D*) (EMF197), *mreC(R292H*) (PR109), *ftsH(V41G*) (EMF199), *mreC(G156D) ftsH(V41G*) (PR111), *mreC(R292H) ftsH(V41G*) (PR110) were incubated in M9 + CAA + glu at 30°C for 24 hours. Cultures were diluted and plated as in **Fig. 1**. **B**. Cultures of the strains listed in (A) were diluted to OD_600_= 0.05 in M9 + CAA + glu and incubated at 30°C until OD_600_= 0.2-0.3. Cultures were gently spun down and resuspended in LB to an OD_600_= 0.025 and incubated at 37°C until OD =0.2-0.3. Cells were fixed and imaged (see methods). Aspect ratios were analyzed using the FIJI plugin MicrobeJ (70). Scale bar = 5 μm. n= 100 cells per group. Statistical significance was determined as in **Fig. 1**. **C**. Cultures of the strains listed in (A) were grown as described in (B) and an immunoblot for LpxC was performed. **D**. Cultures of the strains listed in (A) were grown as described in (B) and analyzed via silver stain for lipid A (top). Samples were normalized to total protein and an immunoblot for RpoA was performed to serve as a loading control.

We chose to further characterize the mechanism of suppression by the *ftsH(V41G*) allele by determining its effect on the cellular concentration of LpxC (**Fig. 2C**) and LPS (**Fig. 2D**). In cells with wild-type FtsH, mutants encoding defective MreC variants had decreased levels of both LpxC (**Fig. 2C**) and LPS (**Fig. 2D**) compared to cells with MreC(WT). The *ftsH(V41G*) allele increased LpxC and LPS levels in all cells regardless of which *mreC* allele they encoded (**Fig. 2C-D**). This change resulted in elevated levels of LPS production in cells with MreC(WT) and an increase in LPS concentration to near normal in cells with the defective MreC variants (**Fig. 2C-D**). We therefore conclude that the *ftsH(V41G*) allele is hypomorphic, leading to reduced LpxC turnover and a rise in LPS levels that compensates for the apparent defect in LPS synthesis of the *mreC* mutants.

To determine whether an increase in LPS synthesis is sufficient to suppress the defective *mreC* alleles, we overexpressed *lpxC* in the mutants (**Fig. 3**). Overproduction of LpxC indeed promoted the growth of *mreC(R292H*) and *mreC(G156D*) mutants on LB and restored an elongated rod-like shape (**Fig. 3**). However, suppression was not as robust as that promoted by the *ftsH(V41G*) allele (**Fig. 2 and 3**), suggesting either that the levels of LPS upon LpxC overproduction were too high and caused mild toxicity or that changes in the turnover of FtsH substrates other than LpxC contribute to the suppressing activity of *ftsH(V41G*). Suppression was dependent on LpxC activity as the overproduction of a catalytically defective LpxC that lacks a degradation signal (designated as ΔC5) (29, 39, 40) failed to promote the elongated growth of cells producing the MreC variants (**Fig. 3**). Notably, overexpression of *lpxC* did not suppress an *mreC* deletion (**Fig. 3**), arguing that partial Rod system activity in the *mreC(R292H*) and *mreC(G156D*) mutants is required to promote rod shape under suppressing conditions. Overall, our results suggest that the growth and shape defects of the *mreC(R292H*) and *mreC(G156D*) mutants is not just due to problems with PG biogenesis. Surprisingly, improper LPS synthesis and OM biogenesis also appear to be contributing factors.

**Figure 3:**
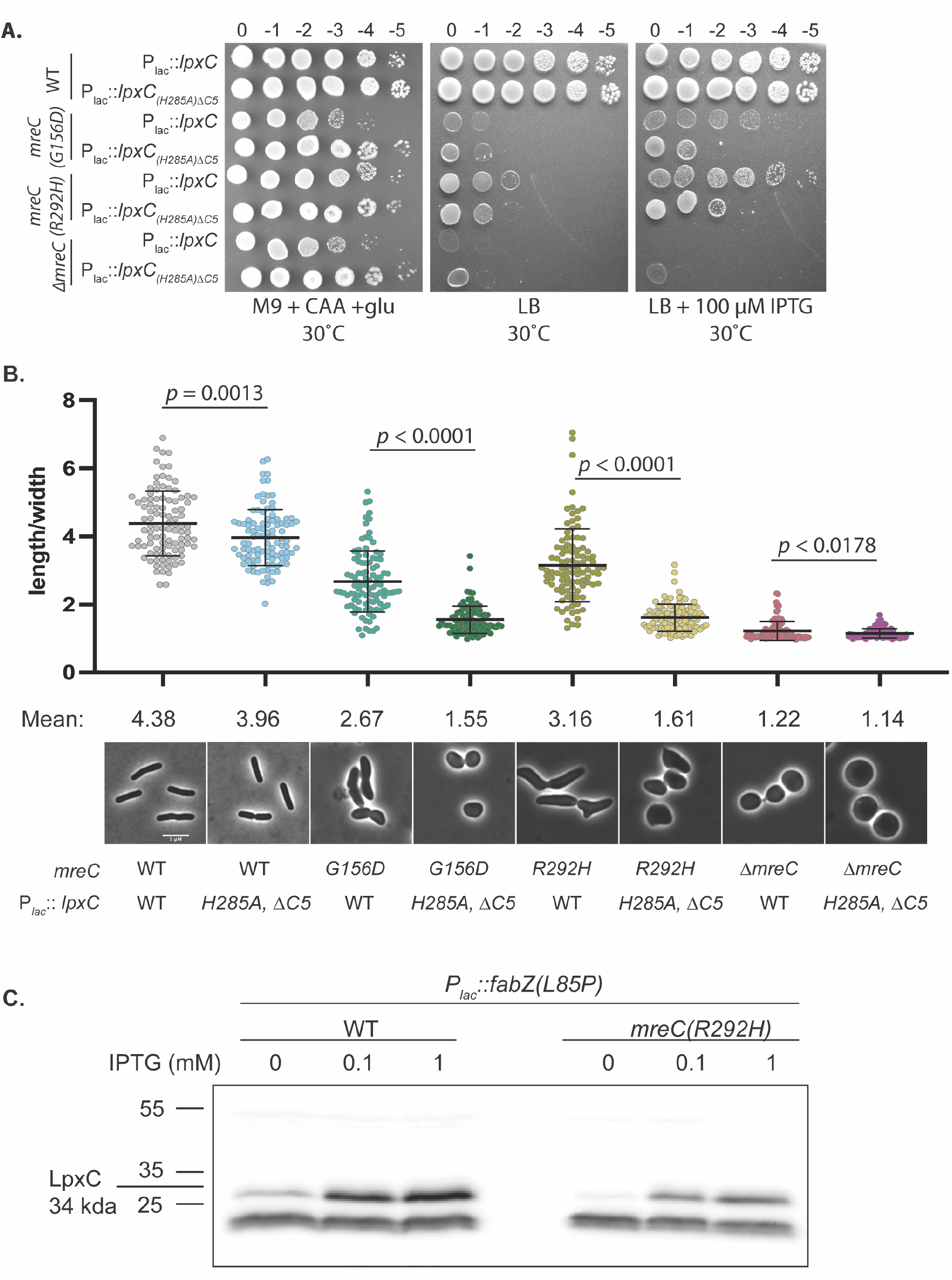
The overexpression of *lpxC* restores growth and partially restores shape to *mreC* hypermorphs. **A**. WT (HC555), *mreC(G156D*) (PR30), *mreC(R292H*) (PR5), and *ΔmreC* (EMF150) expressing WT *lpxC* (pPR111) or *lpxC(H285A)ΔC5* (pPR115) from an IPTG-inducible plasmid were cultured for 24 hours at 30°C in M9 + CAA + glu. Cultures were diluted and plated on the indicated media as in **Fig. 1**. All plates contained chloramphenicol. M9 plates were incubated at 30°C for 40 hours and LB plates were incubated at 30°C for 24 hours. **B**. The strains listed in (A) were grown for 24 hours at 30°C in M9 + CAA + glu + CM. Cultures were diluted to OD_600_= 0.025 in M9 + CAA + glu + CM + 50 μM IPTG and incubated at 30°C until OD_600_= 0.2-0.3. Cells were gently pelleted and resuspended in LB + CM + 50 μM IPTG and grown at 37°C for 1 hour 45 min. Cells were then fixed and imaged (materials and methods). Aspect ratios were analyzed using the FIJI plugin MicrobeJ (70). Scale bar = 5 μm. n= 100 cells per group. Statistical significance was determined as in **Fig. 1**. **C**. Immunoblot for LpxC. Cell lysates of WT (HC555) and *mreC(R292H*) (PR5) cells harboring plasmids expressing *fabZ(L85P*) from an IPTG-inducible promoter (pEMF137) were cultured in M9 + CAA + glu + CM at 30°C for 24 hrs. Cultures were then diluted to OD_600_= 0.025 in M9 + CAA + glu + CM and grown at 30°C until OD_600_= 0.2-0.3. Cells were gently pelleted and resuspended in LB + CM +/- IPTG as indicated and grown at 37°C for 2 hours and were subsequently harvested via centrifugation and processed for immunoblotting.

### mreC mutants remain capable of sensing perturbations to LPS synthesis

One explanation for the decrease in LPS production observed in the *mreC* mutants is that these cells are defective in modulating LpxC stability through the YejM/LapB/FtsH pathway in response to reduced LPS levels (30, 33–38). To test this possibility, we monitored LpxC levels following the overproduction of a hyperactive allele of *fabZ* (3-hydroxy-acyl-[acyl-carrier-protein] dehydratase) (29), an enzyme that functions early in the phospholipid synthesis pathway (41). Overproduction of this enzyme is expected to increase the flux of common precursors into the phospholipid synthesis pathway at the expense of LPS synthesis. Cells harboring the hyperactive *fabZ(L85P*) allele were previously reported to have increased levels of LpxC, presumably due to LpxC stabilization in order to restore balance between the two lipid biosynthesis pathways (29, 42). We found that *mreC(R292H*) cells overexpressing *fabZ(L85P*) had increased levels of LpxC compared to the uninduced controls, and that the magnitude of the increase was comparable to that in WT cells upon induction of the hyperactive *fabZ* allele **(Fig. 3C)**. We observed a similar result when we treated *mreC(R292H*) cells with the LpxC inhibitor CHIR-090 (43, 44), which was also previously shown to promote LpxC stabilization (45) (**SI Fig. 1**). Thus, *mreC* mutant cells remain capable of sensing an acute reduction in LPS synthesis but fail to respond to and correct their chronic deficit in LpxC and LPS levels.

### OM modifications associated with increased stiffness suppress cell shape defects

We reasoned that increasing LPS synthesis could suppress the shape defect of *mreC* mutants either by activating the Rod complex similar to previously characterized suppressors in *rodA* and *mrdA* encoding RodA-PBP2 (10) or by altering the structural properties of the OM. To test the former possibility, we measured the effect of the *ftsH(V41G*) allele on Rod complex activity *in vivo* using a radiolabeling assay. For this assay, a genetic background is used where PG synthesis by the divisome and the aPBPs can be inhibited by SulA production (46–48) and (2-sulfonatoethyl) methanethiosulfonate (MTSES) treatment (49), respectively. Rod system activity can be further isolated by treatment with the PBP2 specific inhibitor mecillinam. This drug blocks the crosslinking activity of PBP2, but the glycosyltransferase activity of RodA remains active, leading to an accumulation of uncrosslinked glycan chains. These uncrosslinked glycans are known to be rapidly degraded by the lytic transglycosylase Slt (49). Thus, the accumulation of nascent PG turnover products during radiolabeling in the presence of mecillinam, MTSES, and SulA can be used as an indirect measure of Rod system activity. Unlike the suppressing RodA and PBP2 variants characterized previously (10) that activate nascent PG turnover product accumulation, the *ftsH(V41G*) allele did not significantly alter Rod complex activity as assessed using the turnover assay **(SI Fig. 2**). Furthermore, the activated PBP2(L61R) variant was found to increase the resistance of cells to the MreB inhibitor A22, another indication of its ability to activate the Rod system. By contrast, overexpression of *lpxC* did not increase resistance to A22 (**SI Fig. 3**). Taken together, these results suggest that hyperactivation of LPS synthesis does not suppress the shape and growth defects of *mreC* mutants by enhancing the PG synthesis activity of the Rod complex.

To investigate whether the mechanical stabilization of the OM is the underlying mechanism by which increased LPS synthesis restores shape to the *mreC* mutants, we sought alternative ways to increase OM stiffness. LPS is composed of three covalently attached units (50). The base glycolipid is called Lipid A. It is modified by a core oligosaccharide that is conserved among Gram-negative organisms. The core is further modified by longer polysaccharide chains called O-antigens, the composition of which varies between species. Laboratory strains of *E. coli* K-12 do not synthesize O-antigen due to an insertion element in *wbbL* (51). However, it was previously reported that restoring O-antigen to the OM dramatically increases its stiffness (3). We therefore asked if re-introducing wild-type *wbbL* to the *mreC* mutants on an arabinose-inducible plasmid could suppress their growth and shape phenotypes like the overexpression of *lpxC* (**Fig. 4A**). Expression of *wbbL* but not a *lacZ* control promoted growth of the *mreC* hypomorphs under the nonpermissive condition (LB, 37°C) and restored their growth as elongated rods (**Fig. 4B-C)**. As we observed with cells overexpressing *lpxC*,overexpressing *wbbL* did not improve the shape or growth defects of *ΔmreC* cells even though they synthesized comparable levels of O-antigen-LPS as the other strains (**Fig. 4D**). Restoring O-antigen synthesis also did not restore shape to cells deleted for *rodZ* (**SI Fig. 4**). Therefore, an intact Rod complex is required to mediate the growth and shape changes in mutant cells with a restored O-antigen. We also investigated if other modifications to the OM can suppress rod system defects (52). We observed the overexpression of *arnT*, which catalyzes the 4-amino-4-deoxyl-L-aminoarabinose (L-Ara4N) modification of lipid A(53, 54), confers a modest but consistent improvement in growth of *mreC(R292H*) mutants under the nonpermissive condition (**SI Fig. 5A**). From these results, we infer that OM stiffening is the likely mechanism by which changes in LPS synthesis or modification restores rod shape to cells with a poorly functioning Rod system.

**Figure 4:**
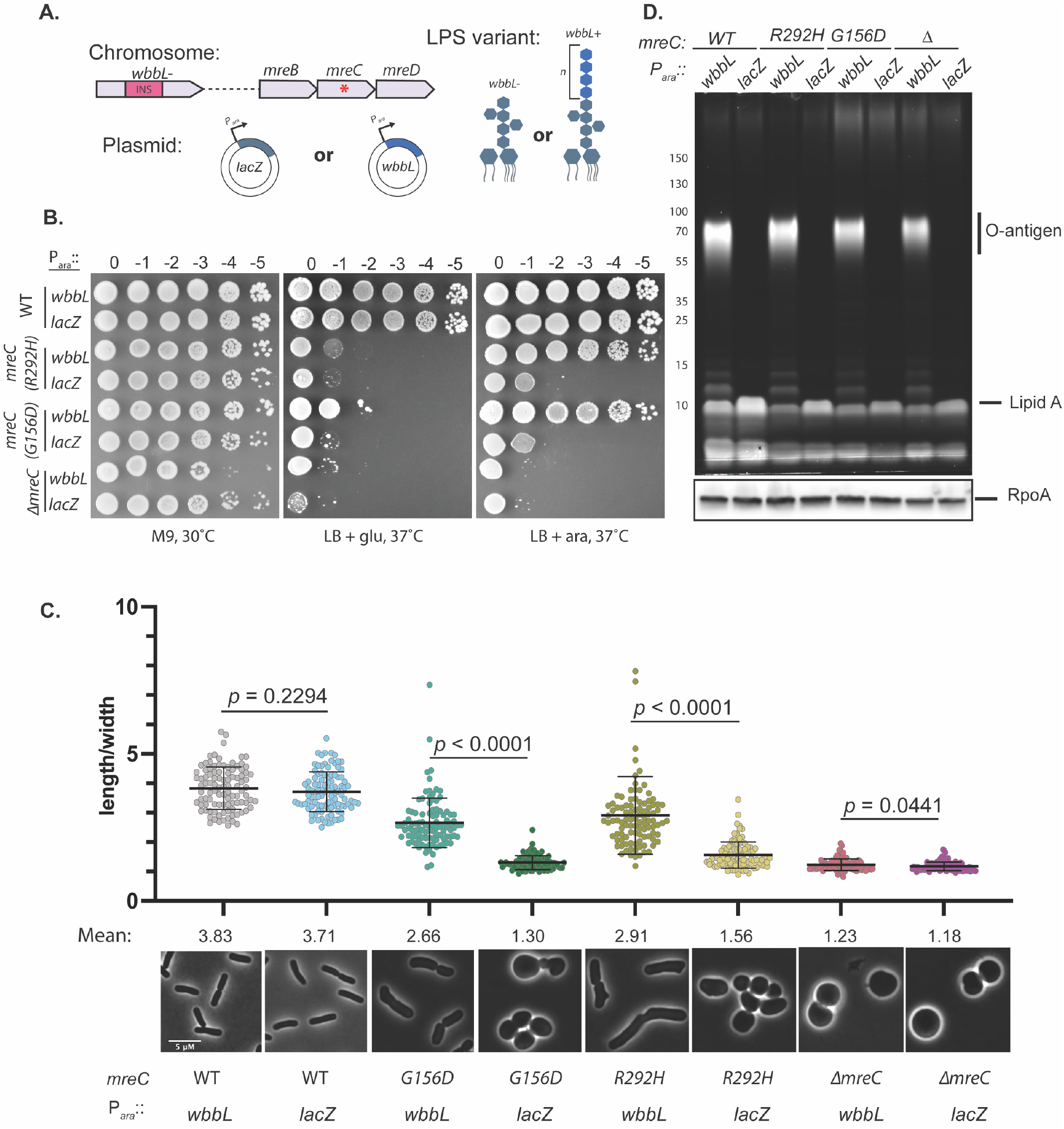
Synthesis of O-antigen-modified LPS suppressed the growth and shape defects of *mreC* hypomorphs. **A**. Schematic of strains. The *wbbL* gene in *E. coli* K-12 is disrupted by an insertion element, preventing the synthesis of O-antigen. *wbbL* is expressed *in trans* from an arabinose (ara)-inducible promoter, restoring O-antigen synthesis. *lacZ* is expressed as a control. **B**. WT (HC555), *mreC(R292H*) (PR5), *mreC(G156D*) (PR30), and *ΔmreC* (EMF150) expressing *wbbL* (pEMF130) or *lacZ*(pEMF134) from an arabinose-inducible promoter were incubated for 24 hours in M9 + CAA + glu + tet at 30°C. Cultures were diluted and plated on the indicated media as in **Fig. 1**. **C**. The strains listed in (A) were grown for 24 hours in M9 + CAA + glu + tet at 30°C and diluted to OD_600_= 0.05 in M9 +CAA + ara + tet for 3 hours at 30°C. After 3 hours, the cultures were gently pelleted and resuspended in LB + tet + ara. Cells were grown for 2 hrs at 37°C. Cells were then fixed and imaged (materials and methods). Aspect ratios were analyzed using the FIJI plugin MicrobeJ (70). Scale bar = 5 μm. n= 100 cells per group. Statistical significance was determined as in **Fig. 1**. **D**. Proemerald Q stain of LPS. The strains listed in (A) were grown as described in (B). Cell lysates were prepared and LPS was analyzed via promerald Q straining.

### OM stiffness and the directional motion of MreB filaments

MreB polymers align along the greatest principal curvature of the cell and are thought to orient the insertion of new PG by the Rod system perpendicular to the long cell axis via a rudder-like mechanism (55). MreB polymers thus promote growth in a rod shape, but they also require rod shape for their proper alignment. Rod-shape is therefore thought to be a self-reinforcing property (21). We reasoned that this rod-shape feedback loop is impaired in the *mreC* mutants because the reduced activity of the Rod system fails to build an envelope robust enough to maintain the beginnings of a cylindrical extrusion that can be elongated into a rod via oriented MreB motion. However, strengthening of the OM in the suppressors may overcome this problem by stabilizing the envelope, allowing a partially functional machine to promote the self-enhancing shape determination process. To test this hypothesis, we wanted to track the motion of a functional MreB-mNeon sandwich fusion (^SW^MreB-mNeon) (7) in *mreC* hypomorphic cells with and without shape-restoring suppressor mutations. Unfortunately, we were unable to construct strains encoding both the *mreC* hypomorphic alleles and the ^*SW*^mreB-mNeon fusion at the native locus because the combination was toxic. Instead, we produced ^SW^MreB-mNeon from the native *mreB* locus that also contained *mreC(WT*) and overexpressed the dominant-negative *mreC(R292H*) allele from a plasmid in cells with or without O-antigen **(Fig. 5A)**. Overexpression of *mreC(R292H*) caused cells lacking O-antigen to form sphere-like cells, but the shape change was not as dramatic as that observed for cells harboring *mreC(R292H*) as the sole copy of the gene at the native locus. As expected, rod shape was maintained in O-antigen positive cells overexpressing *mreC(R292H*).

**Figure 5:**
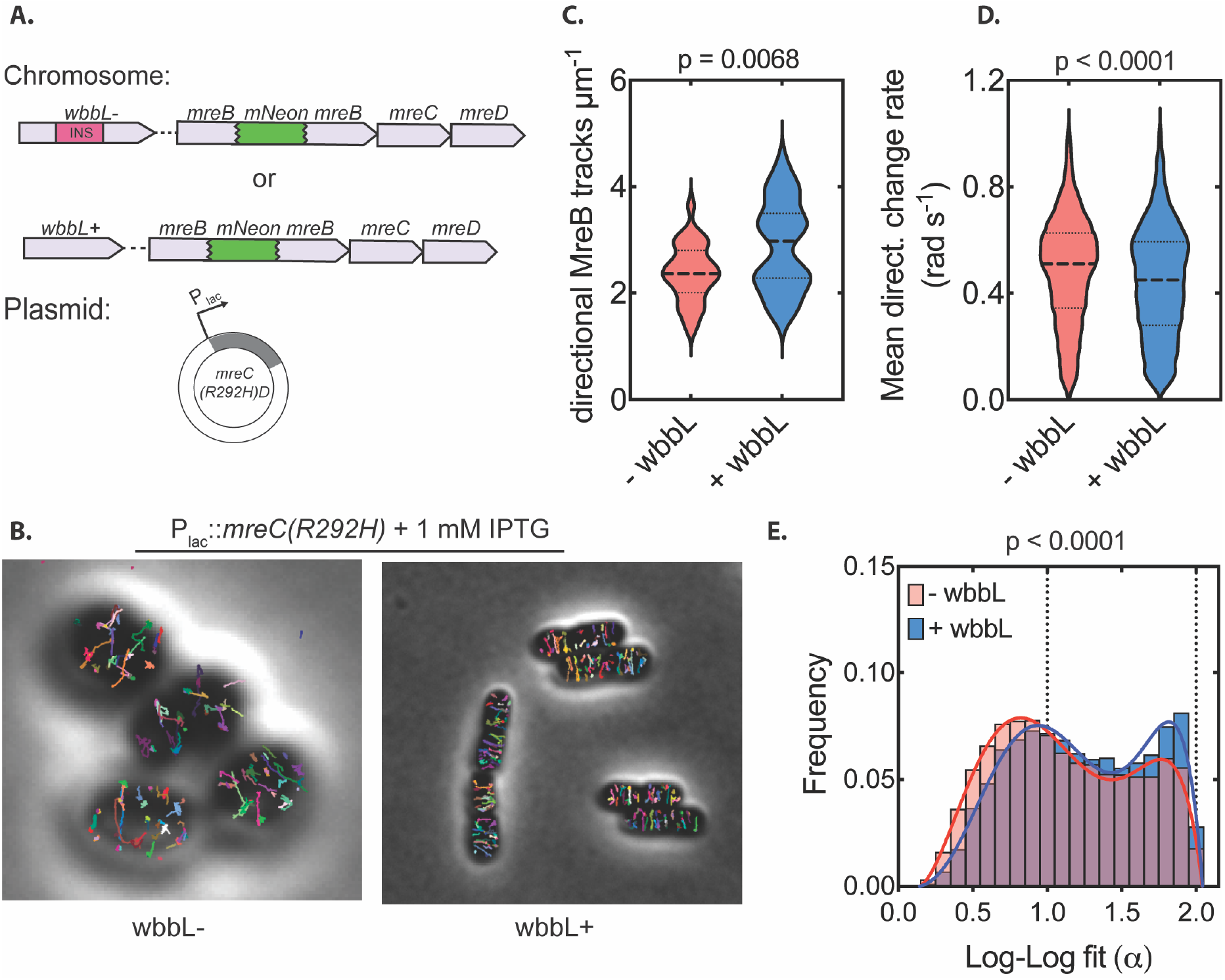
MreB dynamics upon rod system inactivation by *mreC(R292H*) in cells with or without O-antigen. **A**. Schematic of strains. ^*SW*^mreB-mNeon cells harbor either *wbbL*(INS) (AV007) or *wbbL+*(EMF210) at the native chromosomal locus, resulting in cells without or with O-antigen-modified LPS, respectively. *mreC(R292H)D* is expressed in trans from an IPTG-inducible promoter (pMS9). **B**. *wbbL(INS*) (AV007) or *wbbL+* (EMF210) cells expressing *mreC(R292H)D* (pMS9). Individual traces of MreB tracks were mapped using the TrackMate feature of FIJI (72, 75). Each track is indicated in different color. **C**. Violin plot of the number of directional MreB tracks per cell area in cells with (EMF210) and without O-antigen (AV007) expressing *mreC(R292H)D* (pMS9). (n= 30 cells (AV007), n=31 cells (EMF210)) **D**. Violin plot of the mean directional change rate of MreB tracks in *wbbL*(-) and *wbbL* (+) cells (n=10214 tracks (AV007), n= 9162 tracks (EMF210)) **E.** Histogram of the log-log fit (α) values of Individual MreB traces in cells with (EMF210) and without O-antigen (AV007) expressing *mreC(R292H)D* (pMS9). (n=18618 tracks (AV007), n=15070 tracks (EMF210)).

In addition to the differences in shape, the presence of O-antigen also impacted MreB dynamics. Compared to the rod-shaped O-antigen positive cells, cells lacking O-antigen showed a reduction in number of directionally moving particles and those particles that were moving did not appear to have as consistent of an orientation (**Fig. 5A, B, E, supporting movie 1**). Particles in the O-antigen positive cells were also less likely to change direction during imaging than those in the cells lacking O-antigen (**Fig. 5C**). These results argue that the OM contributes to shape determination by providing sufficient envelope stability for MreB directed PG synthesis to be properly oriented and self-reinforcing.

## DISCUSSION

The OM and PG layers of the Gram-negative envelope share numerous connections. Their building blocks are synthesized from common precursors (56–58), and the layers are physically linked by PG binding proteins anchored in the OM (59–61). Additionally, the insertion of beta-barrel proteins in the OM appears to be spatially coordinated with the insertion of new PG material into the mature cell wall matrix (62). Despite these connections, it has only recently been appreciated that the OM plays a role in the mechanical stability of the Gram-negative envelope that rivals that of the cell wall (3, 63). Here, we provide evidence that rather than just stiffening the envelope, the OM also plays a critical role in rod shape determination. Additionally, our genetic analysis uncovered an unexpected connection between LPS synthesis and the activity of the Rod system that elongates the PG matrix, revealing yet another link between the two outermost layers of Gram-negative cells.

A morphogenic role for the OM is inferred from the ability of elevated LPS synthesis or O-antigen modification to restore rod-like shape to cells with a partially defective Rod system. The shape mutants showed a reduced level of LPS and the LPS synthesis enzyme LpxC **(Fig. 2)**. The stiffness of the OM is thought to be mediated by the lateral packing of LPS molecules bridged by Mg^2+^ ions (3). Thus, the OM of the shape defective cells with reduced LPS likely has suboptimal LPS packing and reduced stiffness. Increasing LPS synthesis in these cells by stabilizing LpxC or overproducing it is expected to increase the LPS concentration in the OM of these cells, enhancing lateral interactions between LPS molecules to at least partially restore OM mechanical stability. Similarly, the addition of O-antigen is likely to stiffen the membrane despite suboptimal LPS levels because the extended glycan chains facilitate long distance LPS-LPS interactions.

How does OM stiffening rescue the Rod system defect? We propose that is does so by promoting the oriented-synthesis feedback via which the Rod system generates rod shape (21)(**Fig. 6**). A critical feature of this model of shape determination is that rod shape is self-reinforcing due to the curvature preference of MreB filaments that orients them perpendicular to the long cell axis to guide PG synthesis by the Rod system (55). If the cell wall made by the machinery is not stiff enough to hold the beginnings of a cylindrical shape in the face of turgor pressure, as is likely the case in the *mreC* mutants, then the feedback loop that elongates the cylinder to generate rod-shape cannot be initiated (**Fig. 6**). This problem is encountered in Gram-positive bacteria with defects in wall teichoic acid (WTA) synthesis (55). Much like LPS, these anionic cell wall polymers have been proposed to stiffen the envelope through lateral interactions mediated by bridging Mg^2+^ ions (21). Accordingly, mutants with reduced levels of WTA synthesis can be converted from rods to spheres by removing Mg^2+^ from the medium (55). Moreover, cell shape can be restored to *B. subtills* mutants with a partially defective Rod system by the addition of excess Mg^2+^, which presumably rigidifies the envelope via the WTAs (64, 65). We therefore propose that the LPS of Gram-negative bacteria and WTAs of Gram-positive organisms may function similarly to promote cell shape by providing sufficient envelope rigidity to enable the self-reinforcing orientation of PG synthesis by the Rod system.

**Figure 6:**
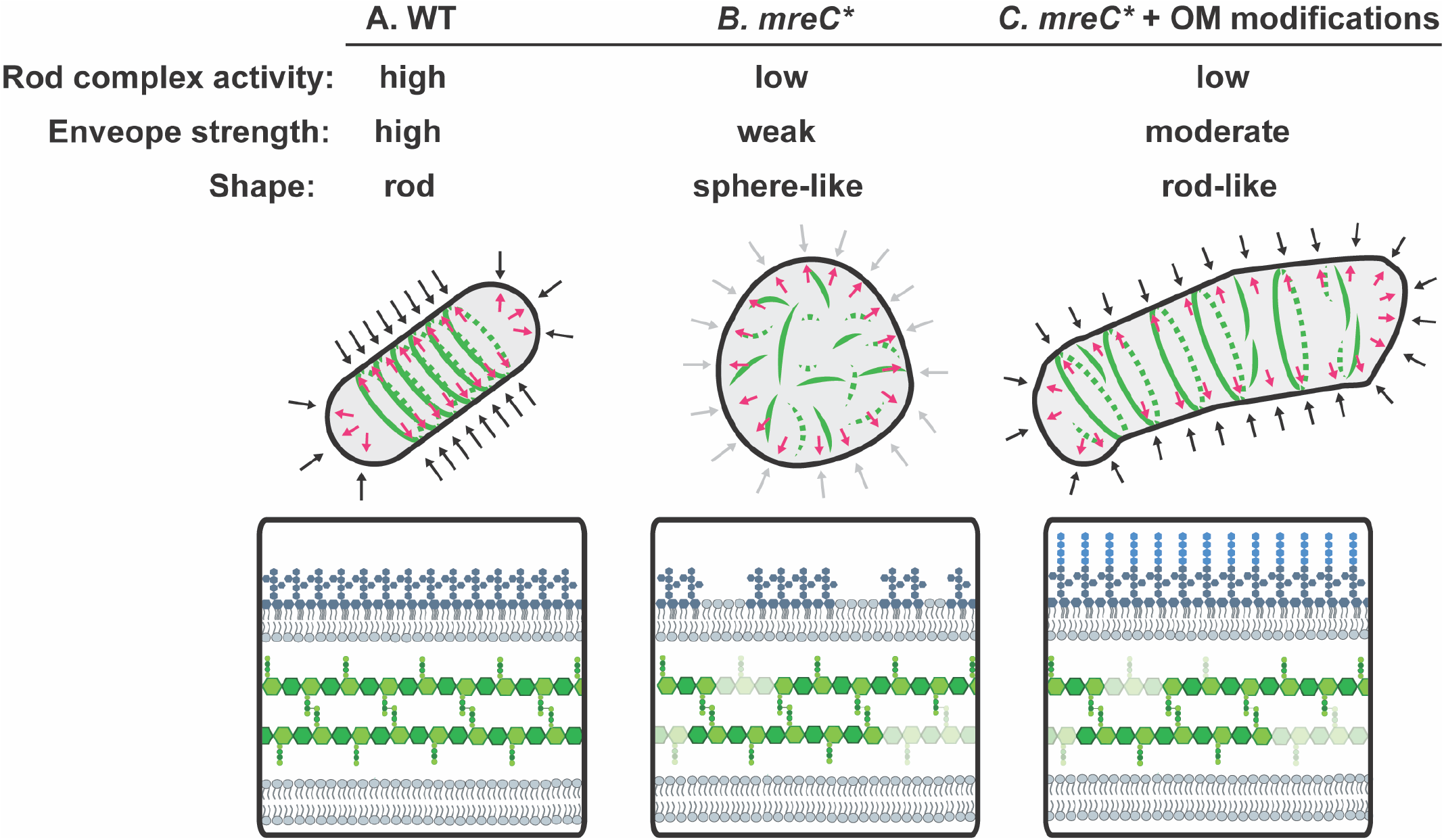
Interventions that strength the outer membrane restore shape to Rod system hypomorphs. **A.** In wildtype cells, the internal turgor pressure of the cell is countered by the combined mechanical strength of the cell wall and the outer membrane. The Rod complex is fully functional and is orientated by MreB, which aligns along the greatest principle curvature to ensure synthesis perpendicular to the long axis of the cell. **B.** In hypomorphic *mreC* mutants (*mreC*\*), the Rod complex is not able to synthesize sufficient peptidoglycan or LPS, weakening the envelope and leading to loss of rod shape. The cells no longer form a clearly defined long axis, causing MreB filaments to misalign. The reduced Rod complex activity in these mutants is therefore not properly oriented. **C.** When the mechanical strength of the outer membrane is increased, the cell envelope is sufficiently able to resist the internal turgor pressure of the cell to allow for the initiation and propagation of a rod shape by allowing MreB and limited PG synthesis by the Rod complex to properly orient.

Given its relevance to antibiotic resistance, the most well-studied role of the OM is as a permeability barrier preventing the entry of bulky and/or hydrophobic drugs. Mutants defective for the Rod system have been known to have a defective OM permeability barrier for many years (11, 66), but the cause of their increased permeability to antibiotics has been unclear. Our results indicate that the problem is likely caused by a reduction in LPS synthesis in the spherical cells. Whether this reflects a direct or indirect connection between Rod system activity and the LPS synthesis and/or transport systems is unclear. However, the *mreC* mutants we studied are still capable of responding to reductions in the flux through the LPS synthesis pathway by stabilizing LpxC **(Fig. 3C, SI Fig 1)**. Thus, the defect does not appear to be at the level of the YejM-LapB-FtsH system that monitors the steady-state level of LPS in the outer leaflet of the IM (30, 33–38). Further investigation of the LPS synthesis defect in cells with reduced Rod system activity may therefore reveal new connections between the biogenesis of the PG and OM layers of the envelope and uncover new ways to compromise the permeability barrier of Gram-negative bacteria to sensitize them to antibiotics.

## METHODS

### Bacterial strains and growth conditions

The strains generated and used in this study are derivatives of MG1655 and cultured in LB (1% tryptone, 0.5% yeast extract, 0.5% NaCl) or minimal (M9) medium (67). Minimal medium was supplemented with 0.2% Casamino Acids and 0.2% glucose (glu) or arabinose (ara) where indicated (see figure legends). Rod system mutants and controls were maintained on M9 + CAA + glu at 30°C unless otherwise indicated. Strains harboring plasmids were grown in the presence of antibiotics at the following concentrations (unless indicated differently in the figure legends): 25 μg/ml chloramphenicol (CM), 25 μg/ml kanamycin (Kan), and 10 μ/ml tetracycline (Tet). All strains, plasmids, and primers used in this study are listed in **SI Tables 2, 3**, and **4**, respectively. For details, please see supplementary text.

### Suppressor analysis

Suppressors were isolated and analyzed as described previously (10).

### Western blots

Cells were pelleted via centrifugation and resuspended in resuspended in water and 2x Laemmli sample buffer (100 mM Tris-HCl, pH 6.8; 2% SDS; 0.1% bromophenol blue; 20% glycerol) at a 1:1 ratio to a final OD_600_ of 20, boiled for 10 minutes, and stored at −80°C. Samples were thawed and sonicated for 1 min twice using a Qsonica tip sonicator with an amplification of 25%. Sample concentration was determined using the Noninterfering (NI) Protein Assay (with bovine serum albumin [BSA] protein standard) (G Biosciences catalog no. 786-005). Samples were run on a 15% polyacrylamide gel (LpxC western blots) or 4–20% Mini-PROTEAN gels (BioRad cat# 4568095) and transferred to a polyvinylidene difluoride (PVDF) membrane. The membrane was rinsed in phosphate-buffered saline containing 0.1% Tween (PBS-T) (10% 10x PBS-T buffer, pH 7.4 [Sigma-Aldrich]) and blocked in 5% milk in PBS-T for 1.5 hours. The membrane was incubated in 1% milk-PBS-T containing rabbit anti-LpxC antibody (a generous gift from the Doerrler lab) or mouse anti-RpoA (anti-*E.coli* RNA polymerase alpha from Biolegend, cat# 663104) diluted 1:10,000. The membranes were incubated at 4°C O/N rocking and then washed 4x with PBS-T at room temperature (1x quickly followed by 3x for 10 min). For LpxC blots, the membrane was incubated in 0.2% milk dissolved in PBS-T with [HRP]-conjugated anti-rabbit IgG (1:40,000 dilution, Rockland cat# 18–8816-33). For RpoA western blots, membranes were incubated with anti-mouse IgG HRP at a dilution of 1:3000 (Thermo Fisher Scientific catalog no. 34577). Membranes were incubated with secondary antibody for two hours and then washed 5x with PBS-T (1x quickly followed by 4x for 10 min per wash). Membranes were developed using the SuperSignal West Pico Plus chemiluminescent substrate (Thermo Fisher Scientific catalog no. 34577) and imaged using the c600 Azure Biosystems platform.

### Detecting LPS using silver stain

Cultures were prepared as described in figure legends. For **Fig. 2**, strains listed in the figure legend were cultured for 24 hours at 30°C in M9 + CAA + glu. Cultures were then diluted to OD_600_= 0.05 and grown at 30°C until OD = 0.2-0.3. Cells were gently pelleted and resuspended in LB (OD_600_= 0.025) and grown at 37°C until OD_600_= 0.2-0.3. Cells were pelleted and resuspended in 1x LDS sample buffer (Invitrogen NP0008) + 4% -mercaptoethanol) to a final OD_600_ of 20. Pellets were boiled for 10 minutes and stored at −80°C. The protein concentration of the samples was measured using the Noninterfering (NI) Protein Assay (with bovine serum albumin [BSA] protein standard) (G Biosciences catalog no. 786-005). RpoA western blots were carried out as described above. For the LPS silver stain, 50 μL of sample was incubated with 1.25 μL of proteinase K (NEB P8107S) for 1 hour at 55°C then 95°C for 10 min. 20 μg (volume equivalent) was resolved on a 4-12% Criterion XT Bis-Tris gel (Bio-Rad 3450124) at 100V for 2 hours. LPS detection via silver stain was performed as described previously (68). First, the gel was fixed overnight in a solution of 200 mL of 40% ethanol and 5% acetic acid. Periodic acid was added to the fixative solution (final concentration of 0.7%). Following a 5 min incubation at room temperature, the gel was washed with 200 mL ultrapure H20 (2x for 30 min, 1x for 1 hour). The gel was then incubated with 150 mL of staining solution (0.018 N NaOH, 0.4% NH4OH, and 0.667% Silver Nitrate) for 10 min. The gel was then washed 3x for 15 min in 200 mL ultrapure H20 and developed in developer solution (0.26 mM Citric Acid pH 3.0, 0.014% formaldehyde). The reaction was stopped by removing the developer and replacing it with 100 mL of 0.5% acetic acid. The gel was imaged using the Bio-Rad ChemiDocTM MP Imaging System.

### Detecting LPS using Pro-Q Emerald 300 lipopolysaccharide gel stain kit

WT (HC555), *mreC(R292H*) (PR5), *mreC(G156D*) (PR30), and *ΔmreC* (EMF150) expressing *wbbL* or *lacZ* from an arabinose-inducible promoter were incubated for 24 hours in M9 + CAA + glu + tet at 30°C and diluted to OD_600_= 0.05 in M9 +CAA + ara + tet for 3 hours at 30°C. After 3 hours, the cultures were gently pelleted and resuspended in LB + ara + tet. Cells were grown for an additional 2 hrs at 37°C. Cells were pelleted and resuspended in 1x LDS sample buffer (Invitrogen NP0008) + 4% 2-mercaptoethanol) to a final OD_600_ of 20, boiled for 10 minutes, and stored at −80°C. The protein concentration of the samples was measured using the Noninterfering (NI) Protein Assay (with bovine serum albumin [BSA] protein standard) (G Biosciences catalog no. 786-005). RpoA western blots were carried out as described above. For the LPS proemeraldQ stain, 50 μL of sample was incubated with 1.25 μL of proteinase K (NEB P8107S) for 1 hr at 55°C then 95°C for 10 min. A normalized volume equivalent to 20 μg total protein in the predigested sample was resolved on a 4-12% Criterion XT Bis-Tris gel (Bio-Rad 3450124) at 100V for 2 hours. The Proemerald Q stain was performed following the manufacturer’s instructions (Pro-Q Emerald 300 lipopolysaccharide gel stain kit-Molecular Probes P20495). The gel was imaged using the Bio-Rad ChemiDocTM MP Imaging System.

### Phase contrast microscopy

Phase contrast micrographs in **Fig. 1, 2, 3, 4, SI 4**, and **SI 5** were all taken using cells fixed in 2.6% in formaldehyde and 0.04% glutaraldehyde. After adding the fixative, cells were incubated at room temperature for 1 hour and stored at 4°C for a maximum of three days. To image, cells were immobilized on agarose pads (2%) on 1 mm glass slides (1.5 coverslips). Micrographs in **Fig. 1** were taken using a Nikon TE2000 inverted microscope using a 1.4 NA Plan Apo Ph3 objective and Nikon Elements Acquisition Software AR 3.2. Micrographs in **Fig. 2** were taken with a Nikon Ti Inverted Microscope using a 1.4 NA Plan Apo 100x Ph3 DM objective and with Nikon Elements 4.30 Acquisition Software.

Micrographs in **Fig. 3, 4, SI 3**, and **SI 4** were taken with a Nikon Ti2-E inverted microscope using a 1.45 NA Plan Apo 100x Ph3 DM objective lens and Nikon Elements 5.2 Acquisition Software. Micrographs were processed using rolling ball transformation (radius = 35 pixels) in FIJI (69) prior to length and width quantification using the microbeJ plugin (70). Aspect ratio was calculated by dividing the length measurements by the width measurements. The data was plotted in Graphpad Prism and statistical analysis of aspect ratio done in GraphPad Prism using a parametric unpaired T test assuming gaussian distribution but not equal standard deviation (Welch’s correction). Images were cropped in FIJI (69).

#### 3H-mDAP physiological radiolabeling

Peptidoglycan turnover was determined as described previously (7, 10, 49). Data was plotted on GraphPad Prism.

### MreB Dynamics

*wbbL(INS*) (AV007) or *wbbL+* (EMF210) cells expressing *mreC(R292H)D* (pMS9) were back diluted from overnight cultures (1:200) and grown in LB + 1 mM IPTG and incubated at 37°C until OD_600_= ~0.4. Cells were then back diluted a second time to OD_600_= 0.05 in LB + 1 mM IPTG and incubated at 37°C until OD_600_= ~0.4. # 1.5 high precision coverslips (Marienfeld) were added to a hydrochloric acid and ethanol and cleaned. Cells were placed onto a 2% (w/v) agarose pad in LB + 1 mM IPTG and imaged at RT on a Nikon Ti inverted microscope equipped with Nikon TIRF Lun-f laser illumination, a Plan Apo 100x, 1.45

NA Ph3 objective lens. Images were recorded using an Andor Zyla 4.2 Plus sCMOS camera and Nikon Elements 4.30 acquisition software. Three-minute timelapse series with an acquisition frame rate of 3s were recorded to capture MreB dynamics and overlayed over a single-frame phase contrast reference image using Fiji (69). Particle tracking was performed as described in Navarro et al. (71). Briefly, MreB tracks were detected in TrackMate v6.0.1(72) using LoG detector (0.3 μm radius) and Kalman filter. To analyze the nature of the displacement of each track, the mean square displacement (*MSD*) was calculated using the MATLAB class msdanalyzer (73). Slopes (*α*) of the individual *MSD* curves were extracted using the Log-log fit of the *MSD* and the delay time *τ*. As the maximum delay time 75% of the track length was used. Only tracks which persisted for longer than 4 timepoints (12s) and with a *R^2^* for log [MSO] versus log [t] above 0.95 were included in the analysis. MreB filaments engaged in active cell wall synthesis are displaced by the action of the enzymatic activities of RodA and PBP2b (2, 7, 17–20, 22, 74) and thus it’s *MSD* curves display slopes of *α* ≈ 2 indicative of a transported particle motion above the rate of Brownian diffusion (*α* ≈ 1) or confined motion (*α* > 1). Mean directional change rate was derived from TrackMate and is defined as a measure of the angle between two succeeding links, averaged over all the links of a track and is reported in radians.

## ACKNOWLEDGEMENTS

We would like to acknowledge all the members of the Bernhardt and Rudner labs for their advice and thoughtful comments throughout the course of this work. We also thank Paula Montero Llopis and the other members of the MicRoN (Microscopy Resources on the North Quad) team at Harvard Medical School for their expertise, support, consultation, and services. We thank Bill Doerrler for the generous gift of the anti-LpxC antibody. We thank Andrew Darwin (NYU) and Teru Ogura (Kumamoto University, Japan) for sharing with us the *fabZ(L85P*) mutant and Natividad Ruiz for sharing the *wbbL+* strain NR2528. This work was supported by the National Institutes of Health (R01 AI083365 and U19 AI158028 to T.G.B.) and Investigator funds from the Howard Hughes Medical Institute (T.G.B.). E.M.F. is supported by the National Science Foundation (NSF) Graduate Research Fellowship award. P.D.A.R. was supported in part by a predoctoral fellowship from the Canadian Institute for Health Research. A.V. is supported by a EMBO long-term postdoctoral fellowship (ALTF_89-2019) and a SNF Postdoc Mobility fellowship (P500PB_203143).

## Supporting Information for

### Supporting text

#### Molecular biology

The polymerase chain reaction (PCR) was carried out using Q5 High fidelity polymerase (New England Biolabs) or GoTaq green master mix (Promega) following manufacturer’s instructions. PCR products were purified using the PCR clean up kit from Qiagen or CWBiosciences. Plasmids were isolated using the Miniprep Kit from Qiagen or the plasmid purification kit from CWBiosciences.

#### Strain construction details

##### PR82

See supplementary table 1 and Methods

##### PR86

See supplementary table 1 and Methods

##### PR88

See supplementary table 1 and Methods

##### EMF196

The *yrdE-kan* allele from strain HC555 was transduced into strain PR103 by P1-mediated transduction. Transductants were selected on LB + kan25 and confirmed for the yrdE-kan allele via PCR.

##### PR103

The *leuU-cat-yhbX* allele from strain PR90 was transduced into MG1655 by P1-mediated transduction. Transductants were selected on LB + CM25.

##### PR90

A chloramphenicol resistance cassette was introduced to strain TB10 between *leuU* and *yhbX*(linked to *ftsH*) by lambda red recombineering. A PCR product amplified from pKD3 was generated using Primers leuU-yhbX_P2_F (ACCTTGAAACGATGGTGCCGGTACGCCTTAGTTATAAATTCATATGAATATCCTCCTTAG) and leuU-yhbX_P1_R (TTGACACAATAAAGTGCCAATTATGTCAGTAGAAGGGAAAGTGTAGGCTGGAGCTGCTTC) and electroporated into strain TB10 following the protocol for strain DY329 described previously(1).

##### EMF197

The *yrdE-kan* marker linked to the *mreC(G156D*) allele from strain PR30/pTB63 was transduced into strain PR103 via P1-mediated transduction. Transductants were selected on M9 + CAA + glu + kan25. The *mreC(G156D*) allele was confirmed via sequencing.

##### EMF199

The *yrdE-kan* allele from strain HC555 was transduced into strain PR104 by P1-mediated transduction. Transductants were selected on LB + kan25 and confirmed for the *yrdE-kan* allele via PCR.

##### PR104

The *leuU-cat-yhbX* allele linked to *ftsH(V41G*) from strain PR96 was transduced into MG1655 via P1-mediated transduction. Transductants were selected on LB + CM25 and confirmed for the *ftsH(V41G*) allele via sequencing.

##### PR96

A chloramphenicol resistance cassette was introduced to strain PR88 between *leuU* and *yhbX*(linked to *ftsH*) by lambda red recombineering with pKD46 plasmid following the protocol described previously (2). A PCR product amplified from pKD3 using primers leuU-yhbX_P2_F (ACCTTGAAACGATGGTGCCGGTACGCCTTAGTTATAAATTCATATGAATATCCTCCTTAG) and leuU-yhbX_P1_R (TTGACACAATAAAGTGCCAATTATGTCAGTAGAAGGGAAAGTGTAGGCTGGAGCTGCTTC) and electroporated into strain PR88/pKD46.

##### PR109

The *yrdE-kan* marker linked to *mreC(R292H*) from strain PR5/pTB63 was transduced into strain PR103 by P1-mediated transduction. Transductants were select on M9 + CAA + glu + kanamycin.

##### PR111

The *yrdE-kan* marker linked to the *mreC(G156D*) allele from strain PR30/pTB63 was transduced into strain PR104 via P1-mediated transduction. Transductants were select on M9 + CAA + glu + kanamycin. The *mreC(G156D*) allele was confirmed via sequencing

##### PR110

The *yrdE-kan* marker linked to *mreC(R292H*) from strain PR5/pTB63 was transduced into strain PR104 by P1-mediated transduction. Transductants were select on M9 + CAA + glu + kanamycin. The *mreC(R292H*) allele was confirmed via sequencing.

##### EMF150

The Δ*mreC::kan* allele from strain MT4/pTB63 was transduced into MG1655 via P1-mediated transduction. Transductants were select on M9 + CAA + glu + kanamycin.

##### AV007

The chloramphenicol resistance cassette in strain JAB593 was cured using pcp20 as described previously (3).

##### EMF210

The *wbbL+* allele linked to a kanamycin resistance cassette from strain NR2528 was transduced into AV007 by P1-mediated transduction. Transductants were selected on LB + kan25 and confirmed via PCR.

##### EMF52(attHKHC859)

The *ftsH(V41G*) allele linked to the *leuU-cat-yhbX* marker was transduced from strain PR96 into strain HC533(attHKHC859) by P1-mediated transduction. Transductants were selected on LB + CM25 and the *ftsH(V41G*) allele was confirmed via sequencing.

##### EMF53(attHKHC859)

The *leuU-cat-yhbX* marker was transduced from strain PR90 into strain HC533(attHKHC859) by P1-mediated transduction. Transductants were selected on LB + CM25 and confirmed via PCR.

##### EMF212

MG1655 x P1(EMF211)

The kanamycin resistance cassette downstream of *wbbL(INS*) was transduced form EMF211 into MG1655 by P1-mediated transduction. Transductants were selected on LB + kan25 and confirmed by PCR.

##### EMF211

A kanamycin resistance cassette introduced to strain TB10 downstream of *wbbL(INS*) by lambda red recombineering. A PCR product amplified from pKD4 was generated using primers wbbL_kan_F (TCGCAACTTTGATCGAATTTCATCAGTTTTTCACCCGTAAGCGATTGTGTAGGCTGGAGC) and wbbL_kan_R (ATAAATAGCTTATCCATGCTTATATGCTTACGGCTTTATACTATTCCGAAGTTCCTATTC) and electroporated into strain TB10 following the protocol for strain DY329 described previously(1).

##### EMF214

The *wbbL+* allele linked to a kanamycin resistance cassette from strain NR2528 was transduced into PR134 by P1-mediated transduction. Transductants were selected on M9 + CAA + glu + kan and confirmed via PCR.

#### Plasmid Construction details

##### pPR112

The nativeRBS_lpxC insert was PCR amplified from E. coli K12 genomic DNA using forward primer LpxC_nativeRBS_XbaI5’(CCCC<u>TCTAGA</u>TAATTTGGCGAGATAATACGATGATC) and reverse primer lpxC_3’truncation_HindIII (TGAT<u>AAGCTT</u>ATTAAGGCGCTTTGAAGGCCAACGG) resulting in an amplified PCR product of the nativeRBS and coding sequencing of lpxC lacking the 5 terminal amino acids. Primers contain xbaI and hindIII restriction sites, respectively. The PCR fragment was cloned into empty vector pPR66 using restriction enzymes xbaI and hindIII.

##### pPR115

pPR112 mutated with quickchange mutagenesis using primer lpxC_quickchange_H265A (TACCGCTTATAAATCCGGTGCTGCACTGAATAACAAACTG)

##### pEMF51

The *nativeRBS_fabZ(L85P*) insert was generated by amplifying the *fabZ* locus from strain EMF63 using primers xbaI_fabZ_F (ATCC<u>TCTAGAT</u>GTCGTTTCTTATATTTTGACAGGAAGAG) and hindlll_fabZ_R (TACC<u>AAGCTT</u>TCAGGCCTCCCGGCTACG). This PCR product was digested with restriction enzymes xbal and hindlll and ligated into pNP140.

##### pEMF137

pEMF51 and pPR66 were digested with restriction enzymes xbal and hindlll-HF. The *nativeRBS_fabZ(L85P*) insert from pEMF51 was ligated into the pPR66 vector.

##### pEMF112

The plasmids pNP146 and pPR111 were digested with restriction enzymes xbal and hindlll-HF. The pNP146 vector backbone was ligated with the *nativeRBS_lpxC* insert from pPR111

##### pEMF131

A PCR product was generated by amplifying the *nativeRBS_ pbp2(L61R)rodA* insert from pPR122 using primers pEMF131_F (CAAA<u>TCTAGA</u>TAAGGGAGCTTTGAGTAG) and pEMF131_R (TGAT<u>AAGCTTA</u>TGCGCACCTCTTACACGCTTTTC). The resulting PCR product was digested with restriction enzymes xbal and hindlll-HF and ligated into the vector backbone of pNP146.

##### pPR122

A PCR product was generated by amplifying the *pbp2(L61R*) allele from genomic DNA from strain PR39 using primers XbaI-pbpA (GCTA<u>TCTAGA</u>TAAGGGAGCTTTGAGTAGAAAACG) and Hindlll-pbpA (GCTA<u>AAGCTT</u>TTTATTCGGATTATCCGTCATG). This PCR product was digested with restriction enzymes xbal and hindlll and ligated into the vector backbone of pHC857.

##### pAF2

A PCR product was generated by amplifying the *arnT* locus from MG1655 gDNA using primers arnT_xbal_F (CCCC<u>TCTAGA</u>TTTAAGAAGGAGATATACATATGAAATCGGTACGTTACCTTATCGG) and arnT_hindlll_R (TGAT<u>AAGCTT</u>ATCATTTGGGACGATACTGAATCAGC) to generate the artificalRBS_arnT fragment which was then digested with restriction enzymes xbal and hindlll and ligated into the pPR66 vector backbone.

##### pEMF130

wbbL amplified from gDNA from strain AAY1using prmers wbbL Xbal-RBS-NdeI5’: (<u>TCTAGA</u>TTAAGAAGGAGATATACATATGGTATATATAATAATCGTTTCCCACGG) and wbbL_Hindlll_Rev (<u>AAGCTT</u>TTACGGGTGAAAAACTGATGAAATTCGATCAAAGTTGCG). The resulting PCR product (xbaI-artificialRBS-wbbL) was cloned into vector pNP146 using restriction enzymes xbal and hindlll.

##### pEMF134

lacZ was amplified from MG1655 gDNA using primers xbal_strongRBS_lacZ (ATCC<u>TCTAGA</u>CTTTAAGAAGGAGATATACCATGACCATGATTACGGATTCACTGG) and hindlll_lacZ_R (TGAT<u>AAGCTT</u>ATTATTTTTGACACCAGACCAACTGGTAATG). The resulting PCR product (xbaI-artificialRBS-lacZ) was cloned into vector pNP146 using restriction enzymes xbal and hindlll. *The artificialRBS indicates the RBS of the F10 gene from T7 bacteriophage

**SI Figure 1:**
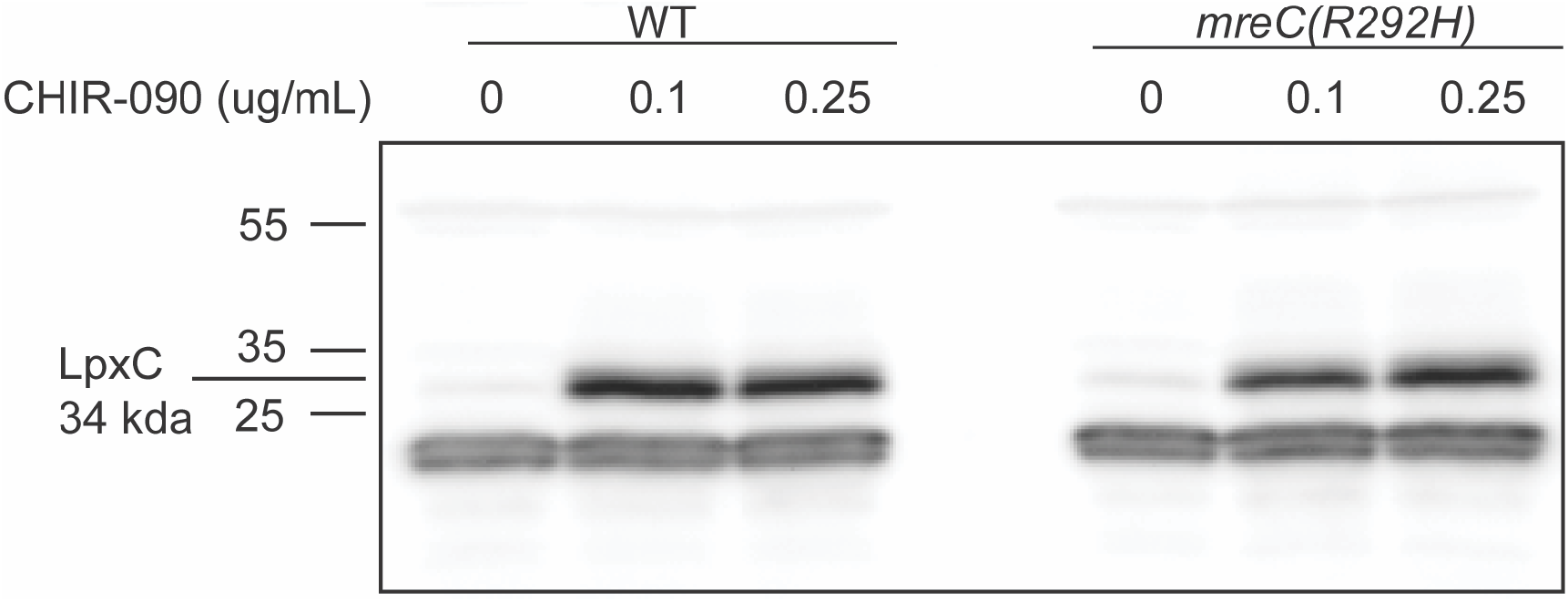
*mreC(R292H*) cells homeostatically regulate levels of LpxC in response to LpxC inhibitor CHIR-090. Immunoblot of LpxC levels in WT (HC555) and *mreC(R292H*) (PR5) cells treated with CHIR-090. Cells were grown for 24 hours at 30°C in M9 + CAA + glu and then back diluted to OD_600_ = 0.05 in M9 + CAA + glu and incubated at 30°C until OD_600_ = 0.4. Cells were gently pelleted and resuspended in LB and grown for one hour at 37°C. CHIR-090 or DMSO was added to the cultures at the indicated concentrations. Cells were incubated for an additional hour at 37°C before cell lysates were harvested for western blot.

**SI Fig. 2:**
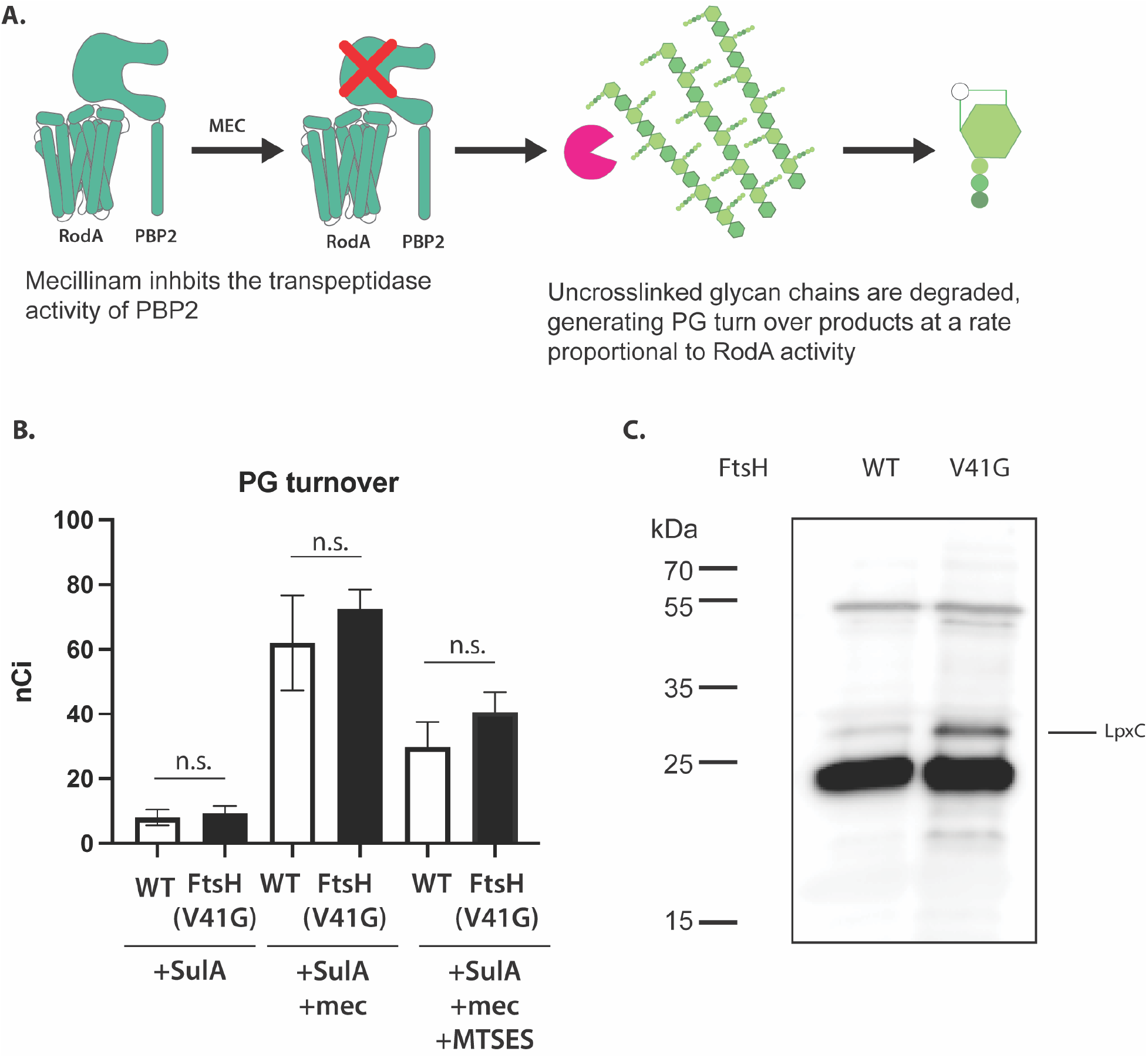
*ftsH(V41G*) does not increase PG synthesis by the Rod complex. **A.** Schematic of the generation of peptidoglycan turnover products (adapted from Rohs et al. 2018(4)). Glycan chains are polymerized by the glycosyltransferase RodA and cross linked into the cell wall matrix by PBP2. Mecillinam blocks the transpeptidases activity of PBP2, leading to the accumulation of uncrosslinked glycan polymers, which are then degraded, generating PG turnover products. These products include a radiolabeled mDAP residue, allowing for detection via HPLC and in-line scintillation counting. **B.** The amount of PG turnover products in WT and FtsH(V41G) cells. SulA blocks divisome activity and MTSES blocks PG synthesis by class A PBPs. Statistical significance was determined using an Unpaired t-test (n.s. indicates not significant). **C.** Immunoblot of LpxC in strains used for radiolabeling assay (see materials and methods).

**SI Figure 3:**
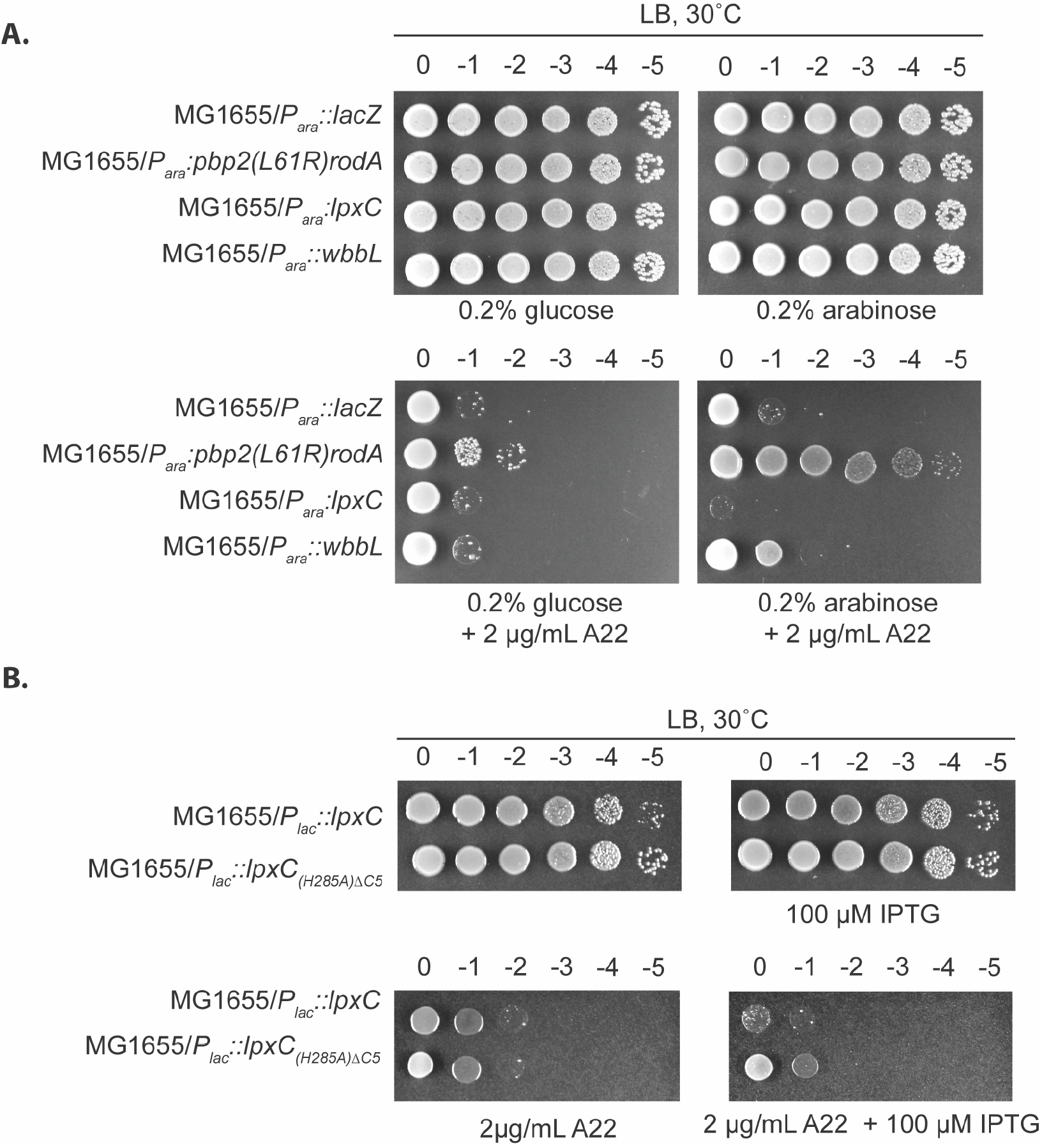
Overexpressing *lpxC* or *wbbL* does not confer A22 resistance. **A.** MG1655 cells harboring arabinose-inducible plasmids expressing *lacZ* (pEMF134), *pbp2(L61R)rodA*(pEMF131) *lpxC* (pEMF112), or *wbbL* (pEMF130) were grown overnight in LB. Cultures were normalized to an OD_600_=1, serially diluted, and spotted on LB + 0.2% glucose or 0.2% arabinose plates with and without 2 μg/mL A22. Plates were incubated at 30°C for 24 hours. **B**. MG1655 cells harboring an IPTG-inducible plasmid expressing *lpxC* (pPR111) or *lpxC(H285A)ΔC5*(pPR115) were grown o/n at 30°C in LB. Cultures were normalized to an OD_600_=1, serially diluted, and spotted on LB plates with and without 100 μM IPTG and 2 μg/mL A22. Plates were incubated at 30°C for 24 hours.

**SI Figure 4:**
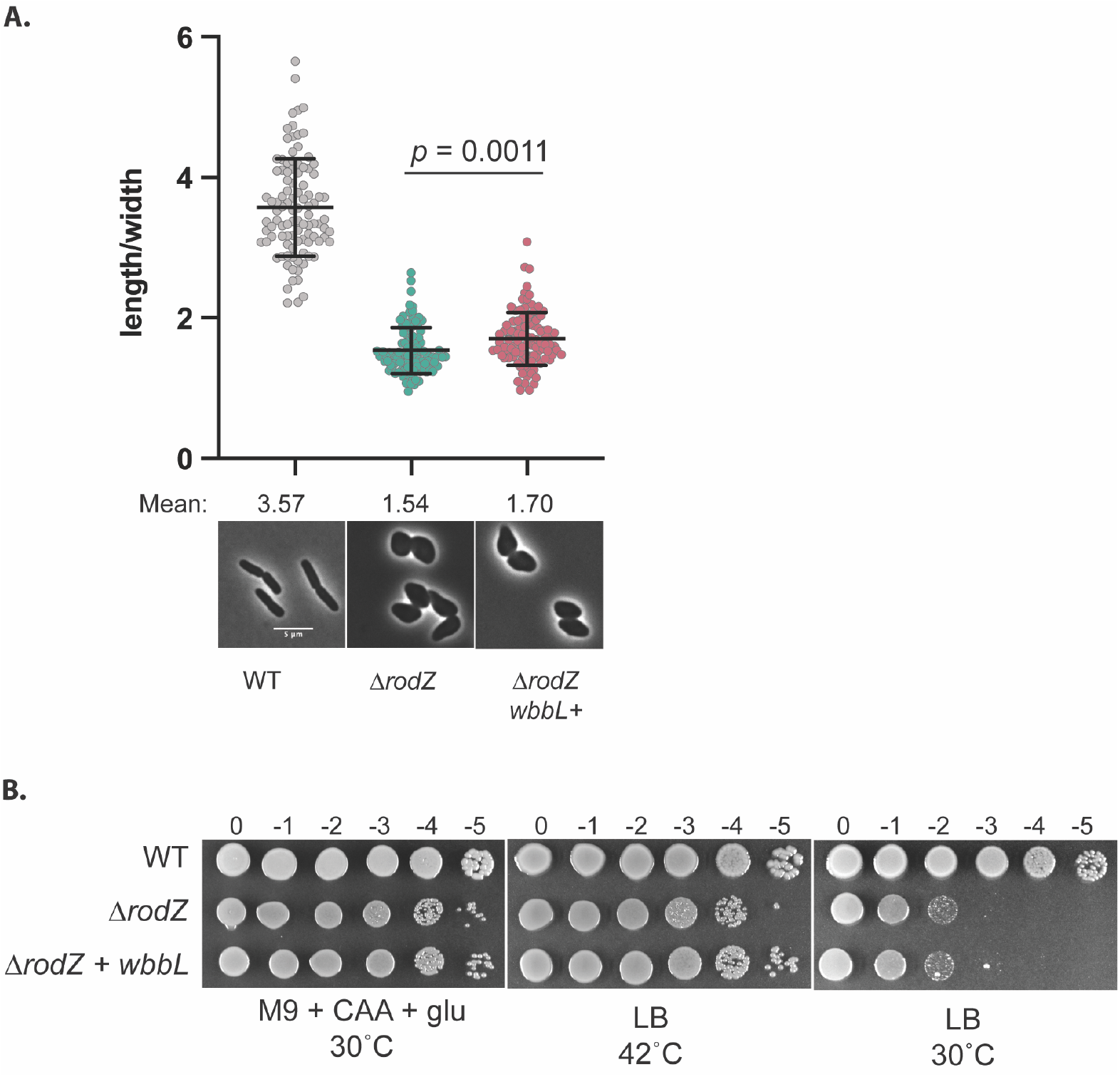
The overexpression of *wbbL* does not ameliorate the growth or shape defects of *ΔrodZ* cells. **A.** WT (EMF212), Δr*odZ* (PR134), and *ΔrodZ wbbL+* (EMF214) were cultures O/N in LB at 37°C. Cultures were diluted at a ratio of 1:200 in LB and grown at 30°C until OD_600_ = 0.3. Cells were then fixed and imaged. Aspect ratios were analyzed using the FIJI plugin MicrobeJ (5). Scale bar = 5 μm. n= 100 cells per group. Statistical significance determined using an Unpaired t test with Welch’s correction (not assuming equal SDs). **B.** The strains listed in (A) were cultures in LB at 37°C overnight. Cultures were normalized to OD_600_=1, serially diluted, and spotted on M9 + CAA + glu or LB plates. LB plates were incubated for 16 hours and M9 + CAA + glu plates were incubated for 48h hours.

**SI Figure 5:**
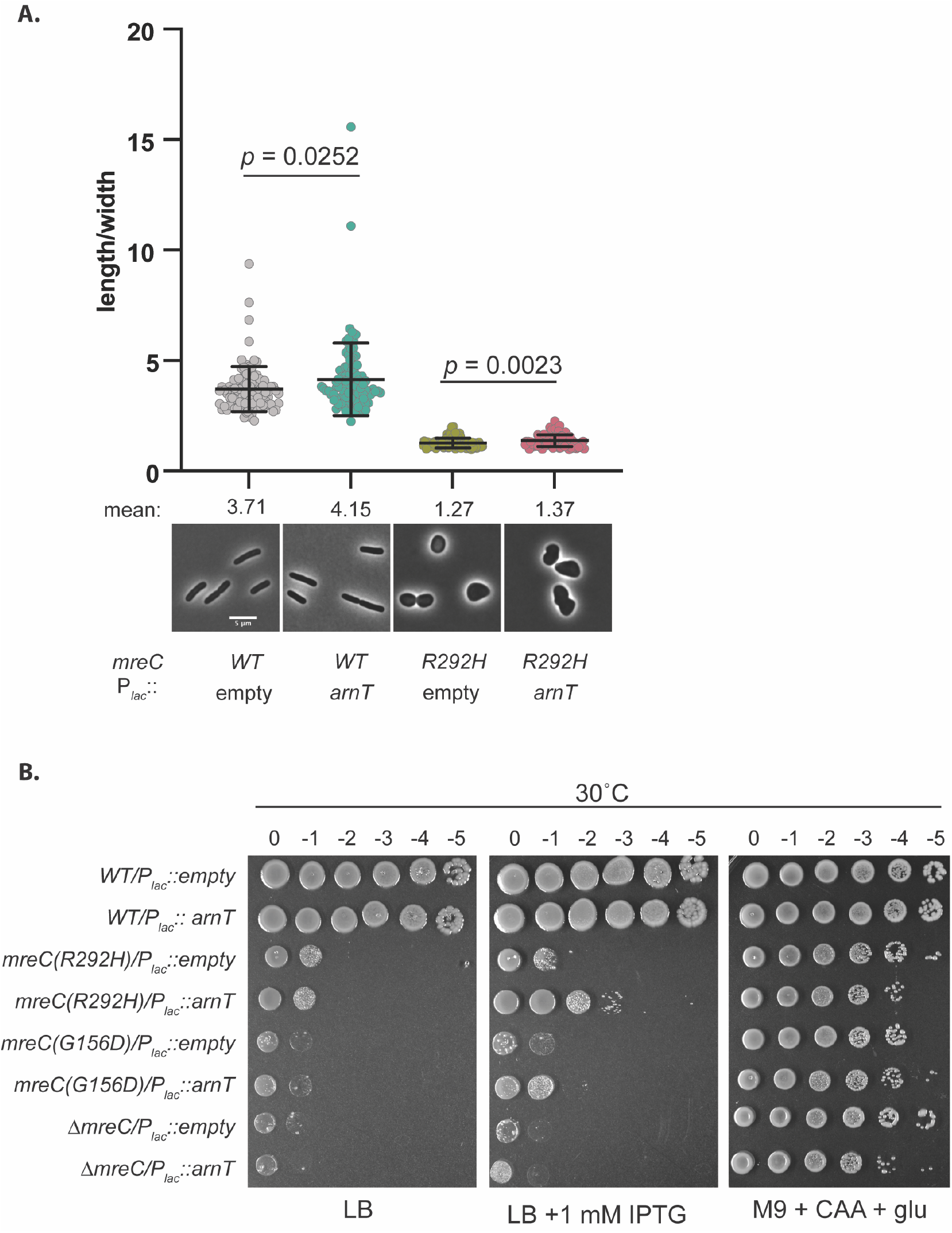
The overexpression of *arnT* partially ameliorates the growth defect of *mreC(R292H*). **A.** WT (HC555) or *mreC(R292H*) (PR5) harboring either the empty vector (pPR66) or an IPTG-inducible plasmid expressing *arnT* (pAF2) were grown for 24 hours at 30°C in M9 + CAA + glu. Cells were diluted to OD_600_= 0.05 in M9 + CAA + glu + 1mM IPTG and cultured at 30°C until OD_600_=0.2-03. Cells were then gently pelleted and resuspended in LB + 1mM IPTG and grown at 37°C until OD_600_ = 0.3. Cells were then fixed and imaged. Aspect ratios were analyzed using the FIJI plugin MicrobeJ (5). Scale bar = 5 μm. n= 100 cells per group. Statistical significance determined using an Unpaired t test with Welch’s correction (not assuming equal SDs). **B.** WT (HC555) or mreC(R292H) (PR5) harboring either the empty vector (pPR66) or an IPTG-inducible plasmid expressing *arnT* (pAF2) were grown for 24 hours at 30°C in M9 + CAA + glu. The overnight cultures were normalized to a OD_600_=1, serially diluted, and spotted LB, LB + 1mM IPTG, and M9 + CAA + glu plates. LB plates were incubated at 30°C for 24 hours and the M9 plates were incubated at 30°C for 48 hours.

**SI Movie 1: Time lapse of MreB in *wbbL(INS*) (AV007) and *wbbL (+*) (EMF210) cells expressing *mreC(R292H)D* (pMS9) described in Fig. 5**. Three-minute timelapse series with an acquisition frame rate of 3s were recorded to capture MreB dynamics. TOP: ^*SW*^mreB-mNeon overlayed over a single-frame phase contrast reference image. BOTTOM: Examples of MreB tracks identified using TrackMate (6, 7). Scale Bar = 2 μM.

**Table S1:** Suppressors of *mreC(R292H*) and *mreC(G156D*)

| Suppressor | Background | Selection strategy | description |
| --- | --- | --- | --- |
| *ftsH(V41G)* | *mreC(G156D)* | Spontaneous suppressors, LB + 1% SDS at 30°C | Inner membrane zinc-dependent metalloprotease that regulates the degradation of UDP-3-O-acyl-N-acetylglucosamine deacetylase (LpxC)(8, 9), the enzyme that catalyzes the first committed step in LPS synthesis (10, 11). |
| *ftsH(F37V)* | *mreC(R292H)* | Spontaneous suppressors, LB at 30°C |  |
| *lapB(Δ379-389)* | *mreC(G156D)* | Spontaneous suppressors, LB + 1% SDS at 30°C | Lipopolysaccharide assembly protein B, mediates LpxC degradation by FtsH (12-14) |

**Table S2:**
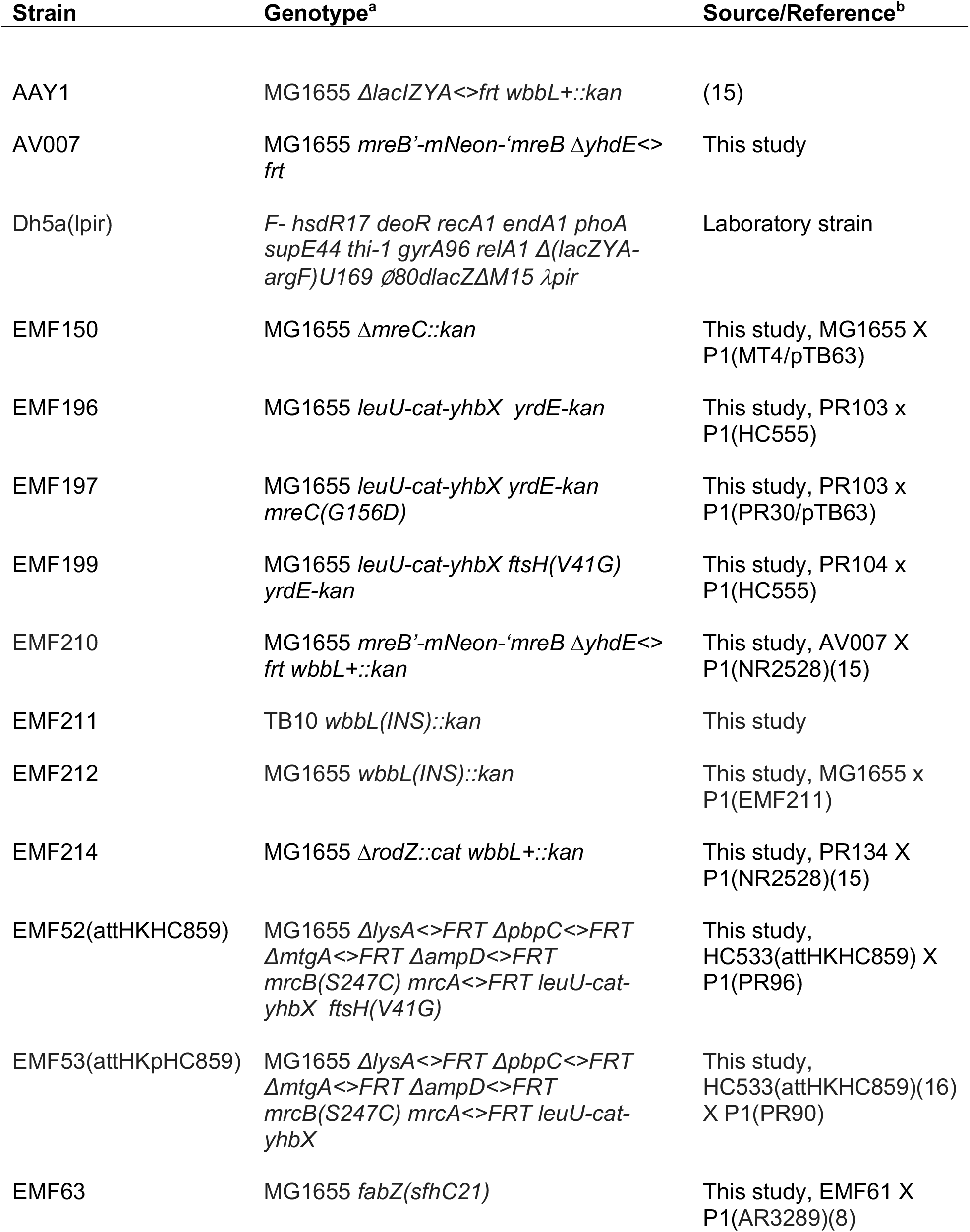
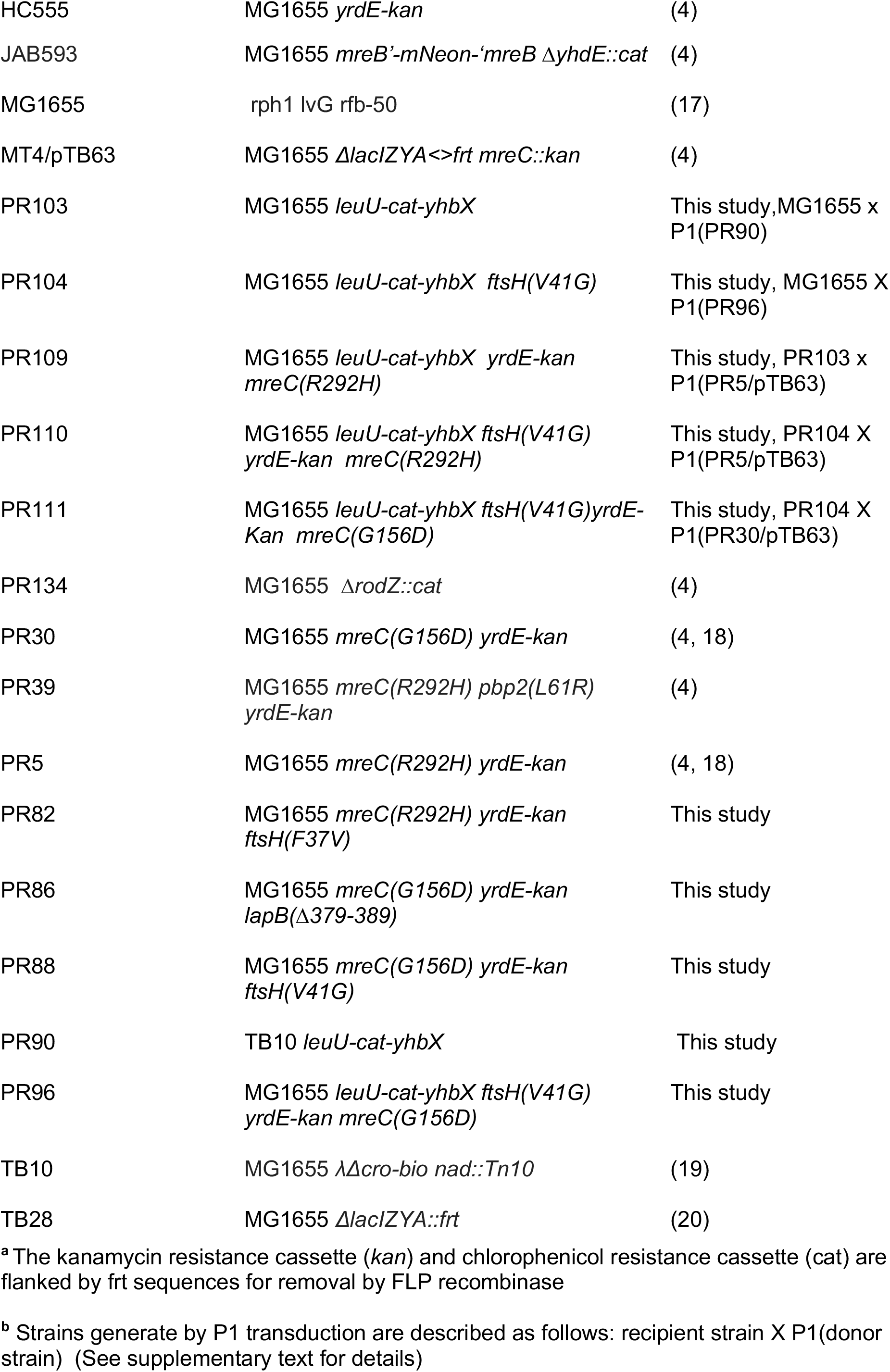
Strains used in this study.

**Table S3:** Plasmids used in this study.

| Plasmid | Relevant features<sup>a</sup> | Origin | Reference/source |
| --- | --- | --- | --- |
| pAF2 | CM<sup>R</sup>, P<sub>lac</sub>::*artificialRBS_arnT* | pBR/colE1 | This study |
| pcp20 | CM<sup>R</sup>, Amp<sup>R</sup>, FLP+, lambda cl857+ | pSC101 | (3) |
| pEMF112 | Tet<sup>R</sup>, P<sub>ara</sub>::*nativeRBS_lpxC* | pBR/colE1 | This study |
| pEMF130 | Tet<sup>R</sup>, P<sub>ara</sub>::*artificialRBS_wbbL* | pBR/colE1 | This study |
| pEMF131 | Tet<sup>R</sup>, P<sub>ara</sub>::*nativeRBS_pbp2(L61R)rodA* | pBR/colE1 | This study |
| pEMF134 | Tet<sup>R</sup>, P<sub>ara</sub>::*artificialRBS_lacZ* | pBR/colE1 | This study |
| pEMF137 | CM<sup>R</sup>, P<sub>lac</sub>:: *nativeRBS_fabZ(L85P)* | pBR/colE1 | This study |
| pEMF51 | Tet<sup>R</sup>, P<sub>ara</sub>:: *nativeRBS_fabZ(L85P)* | pACYC | This study |
| pHC857 | CM<sup>R</sup>, P<sub>lac</sub>::*nativeRBS_pbpA-rodA* | pBR/colE1 | (16) |
| pHC859 | Tet<sup>R</sup>, P<sub>tac</sub>::*sulA* | R6K | (16) |
| pKD46 | Amp<sup>R</sup>, P<sub>ara</sub>::lambda red genes for recombineering | pSC101 | (2) |
| pMS9 | CM<sup>R</sup>, P<sub>lac</sub>::*nativeRBS_mreC(R292H)mreD* | pBR/colE1 | (18) |
| pNP140 | Tet<sup>R</sup>, P<sub>ara</sub>:: *sulA* | pACYC | This study |
| pNP146 | Tet<sup>R</sup>, P<sub>ara</sub>::*sulA* | pBR/colE1 | (21) |
| pPR111 | CM<sup>R</sup>, P<sub>lac</sub>::*nativeRBS_lpxC* | pBR/colE1 | (22) |
| pPR112 | CM<sup>R</sup>, P<sub>lac</sub>::*nativeRBS_lpxCΔC5* | pBR/colE1 | This study |
| pPR115 | CM<sup>R</sup>, P<sub>lac</sub>::*nativeRBS (H285A)_lpxC* | pBR/colE1 | This study |
| pPR122 | CM<sup>R</sup>, P<sub>lac</sub>::*nativeRBS_pbpA(L61R)rodA* | pBR/colE1 | This study |
| pPR66 | CM<sup>R</sup>, P<sub>lac</sub>::empty | pBR/colE1 | (22) |
| pTB63 | Tet<sup>R</sup>, P<sub>native</sub>::*ftsQAZ* | pSC101 | (23) |

<sup>a</sup> P<sub>lac</sub> and P<sub>tac</sub> refer to the lactose promoters and P<sub>ara</sub> refers to the arabinose promoter. The artificialRBS indicates the RBS of the Φ10 gene from T7 bacteriophage.

**Table S4:**
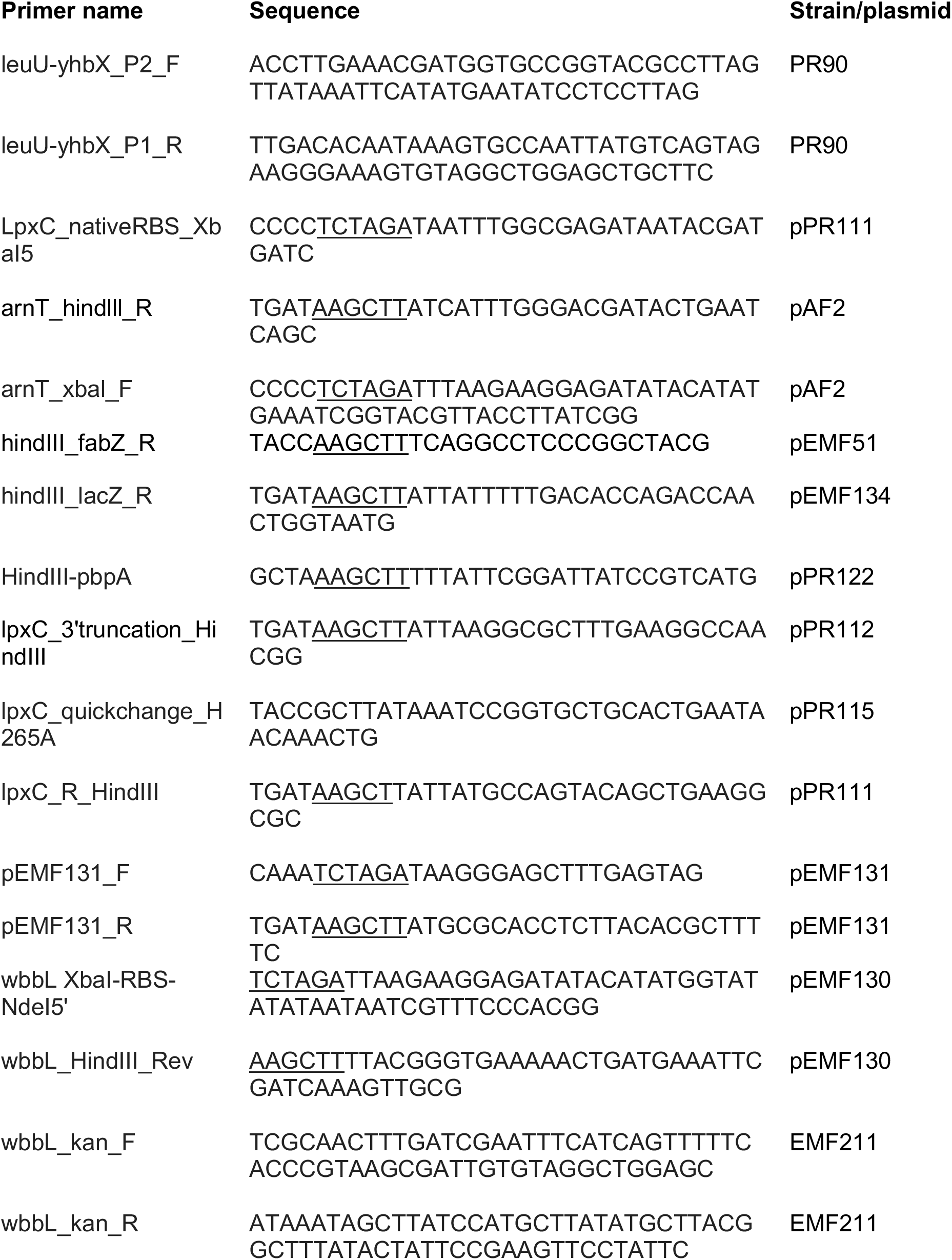
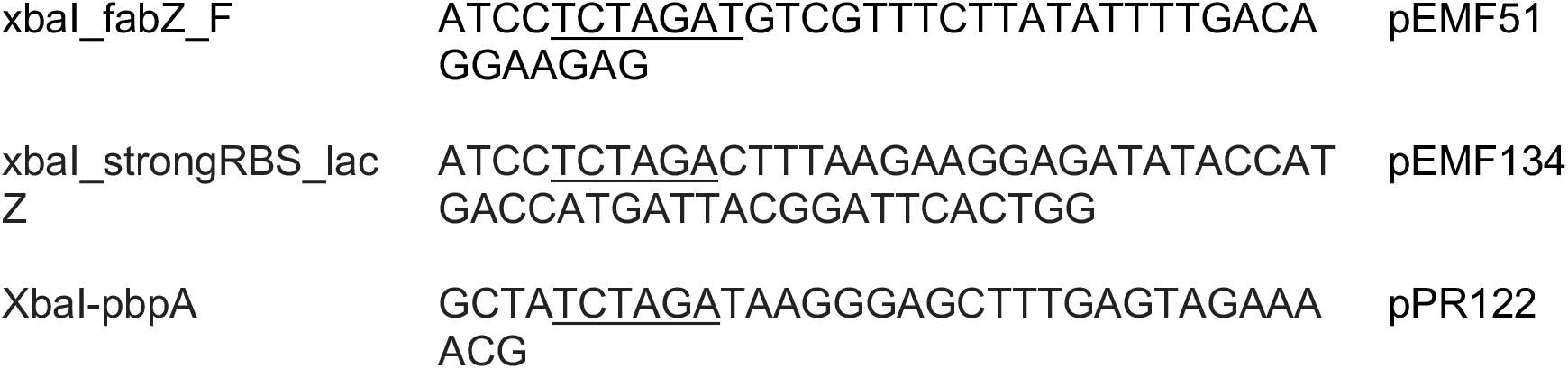
Primers used in this study.

